# Evidence that Dmrta2 acts as a transcriptional repressor of *Pax6* in murine cortical progenitors and identification of a mutation crucial for DNA recognition associated with microcephaly in human

**DOI:** 10.1101/2024.09.20.614077

**Authors:** Xueyi Shen, Jithu Anirudhan, Ambrin Fatima, Tünde Szemes, Marc Keruzore, Estelle Plant, Alba Sabaté San José, Sadia Kricha, Louis-Paul Delhaye, Bilal Ahmad Mian, Lubaba Bintee Khalid, Farhan Ali, Hijab Zahra, Asmat Ali, Mathias Toft, Marc Dieu, Carine Van Lint, Younes Achouri, Patricia Renard, Zafar Iqbal, Eric Bellefroid

## Abstract

Dmrta2 (also designated Dmrt5) is a transcriptional regulator expressed in cortical progenitors in a caudomedial^high^/rostrolateral^low^ gradient with important roles at different steps of cortical development. Dmrta2 has been suggested to act in cortex development mainly by differential suppression of Pax6 and other homeobox transcription factors such as the ventral telencephalic regulator Gsx2, which remains to be fully demonstrated. Here we have addressed the epistatic relation between *Pax6* and *Dmrta2* by comparing phenotypes in mutant embryos or embryos overexpressing both genes in various allelic combinations. We showed that Dmrta2 cooperates with Pax6 in the maintenance of cortical identity in dorsal telencephalic progenitors and that it acts as a transcriptional repressor of *Pax6* to control cortical patterning. Mechanistically, we show that in P19 cells, Dmrta2 can act as a DNA-binding dependent repressor on the *Pax6 E60* enhancer and that a point mutation that affects its DNA binding properties leads to agenesis of the corpus callosum, pachygyria, and the absence of the cingulate gyrus. Finally, we provide evidence that Dmrta2 binds to the Zfp423 zinc finger protein and that it enhances its ability to recruit the NurD repressor complex. Together, our results highlight the importance and conserved function of Dmrta2 in cortical development and provide novel insights into its mechanism of action.

**SIGNIFICANCE STATEMENT:** Corticogenesis is controlled by an array of transcription factors that coordinate neural progenitor self-renewal and differentiation to generate correct cortical cell number and diversity. However, how this complex array of transcription factors works in concert to regulate this delicate process remains largely unknown. Here we provide important insights into the mechanism of action of Dmrta2 by demonstrating that it cooperates with the transcription factor *Pax6* to define the pallium-subpallium boundary and that it acts by repressing it, likely via the recruitment of Zfp423 and the NurD repressor complex, to control cortical patterning. Our data also reveal that a point mutation that affects its DNA binding causes cortical abnormalities in human, further highlighting its importance in cortex development.

## INTRODUCTION

Balancing neural progenitor self-renewal and differentiation is essential for generating cells in correct numbers and diverse types during neural development. During cerebral cortex development, neurogenesis is tightly regulated by a complex array of transcription factors (TFs). Many of these TFs are expressed in graded patterns along the rostral/caudal (R/C) and dorsal/ventral (D/V) axes of the developing ventricular zone (VZ) of the pallium (dorsal telencephalon) under the control of signals produced by localized signaling centers located at the periphery of the cortical primordium (Cadwell et al., 2019; Tole and Hébert, 2020; Ypsilanti et al., 2021). How this host of TFs expressed in gradients in progenitors orchestrate together their proliferation and differentiation to build the cerebral cortex remains today an important unanswered question.

Among the genes coding for cortical TFs is the zinc finger *Dmrta2* gene, also named *Dmrt5*, which is expressed in a caudomedial^high^/ rostrolateral^low^ gradient in the developing cortical VZ. In mice lacking *Dmrta2*, the telencephalic vesicle size is decreased due to premature differentiation of progenitors (Young et al., 2017). The hippocampus and the cortical hem, one of the major telencephalic patterning centers, and the caudal neocortical area are strongly reduced (Konno et al., 2012; Saulnier et al., 2013). The additional knock-out of the related *Dmrt3* gene in mice lacking *Dmrta2* leads to a more severe phenotype with cortical progenitors expressing ventral telencephalic markers such as *Gsx2* (Desmaris et al., 2018). Conditional deletion of *Dmrta2* leaving cortical hem intact also results in a similar phenotype, suggesting that *Dmrta2* controls the patterning of the cortex independently of its influence on the hem. Conversely, in *Dmrta2* overexpressing mice, the caudal area expands while rostral ones are reduced (De Clercq et al., 2018).

Mechanistically, Dmrta2 has been shown to bind to telencephalic enhancers of the *Pax6* and *Gsx2* loci and repress their expression (Konno et al., 2019). While both *Dmrta2* and *Pax6* contribute to telencephalic dorsoventral patterning by repressing ventral telencephalic-specific genes, *Pax6* within cortical progenitors regulates in an opposite manner to *Dmrta2* patterning and arealization of the neocortex (Ypsilanti and Rubenstein, 2016). Although *Pax6* effects are context and dose-dependent, it can promote neuronal differentiation while *Dmrta2* promotes the maintenance of neural progenitors during neurogenesis (Manuel et al., 2015; Young et al., 2017). Therefore, it has been suggested that *Dmrta2* acts through the repression of *Gsx2* to define the dorsal telencephalic compartment and through the repression of *Pax6* to determine positional information within cortical progenitors (Konno et al., 2019). First evidence for the hypothesis that *Dmrta2* acts through *Pax6* in cortex development has been obtained in *Dmrta2* loss of function experiments using electroporated siRNA, in the context of its opposite function to *Pax6* in the control of the differentiation of progenitors. Results obtained showed that the double knock-down of *Dmrt3* and *Dmrta2* upregulates the expression of proneural genes such as *Neurog2* or *Neurod1*, that act as the primary initiators of neuronal differentiation and are directly activated by *Pax6*, while the additional knockdown of *Pax6* in *Dmrt3*; *Dmrta2* double knock-down rescued the neurogenic phenotype (Konno et al., 2019).

In this study, we compared phenotypes of embryos mutant for both *Dmrta2* and *Pax6* in various allelic conditions and double conditional transgenics overexpressing both *Dmrta2* and *Pax6* with transgenics overexpressing only *Dmrta2* to understand how *Dmrta2* and *Pax6* function together in cortex development. We then performed reporter assays in P19 teratocarcinoma cells and rapid immunoprecipitation followed by mass spectrometry of endogenous protein (RIME) experiments to approach the mechanism of action of Dmrta2. We also examined in human a novel case of microcephaly with a point mutation in *DMRTA2*. Together, our results demonstrate the importance of Dmrta2 and Pax6 interactions in the positional information of cortical progenitors and provide first insights into its mechanism of action.

## MATERIALS AND METHODS

### Animal lines and genotyping

All animal experiments were performed with approval from the ethics committee of the (Author University) and conformed to guidelines on the ethical care and use of animals. The midday of the day of vaginal plug discovery was termed embryonic day 0.5 (E0.5), and the first 24 hours (h) after birth were P0. Animal care was in accordance with institutional guidelines and the National Institutes of Health policies. Mice were provided *ad libitum* with standard mouse lab pellet food and water and housed at room temperature (RT) with a 12h light/dark cycle.

*Dmrta2^+/-^* and *Dmrta2^Tg/+^* (De Clercq et al., 2018), *Pax6^sey/+^* (Hill et al., 1991), and *BAT-gal* (Maretto et al., 2003) mice were maintained in a C57Bl/6 background. *Pax6^Tg/+^* (Berger et al., 2007) were in an FVB/N background. The *Dmrta2* knock-in (*Dmrta2^2xHA^*) mouse line was generated using CRISPR/cas9 editing. A crRNA was designed using CRISPRdirect (http://crispr.dbcls.jp/) with the intent to produce a double-strand break in the second exon of D*mrta2*, 5’ from the ATG codon. 2.4 pmol/µl of annealed crRNA (10nmol, Integrated DNA Technologies, 5’-TCCAGCCATGGAGCTGCGCT-3’) and the tracrRNA (Integrated DNA Technologies, catalog #1072534) were injected with 100 ng/μl of recombinant Cas9 protein (Integrated DNA Technologies) and 10 ng/μl of a single single-stranded DNA oligonucleotide template with homology arms to the targeted region and containing a 2XHA sequence (Integrated DNA Technologies, Megamer® ssDNA Fragment: 5’-CTTAAGCCTAGTGACTGTATTACCCTGACTCAGGTATCCCCACTTTCCCCACGCTA GG<u>TCCAGCC</u>**ATG**GAGTATCCCTATGACGTCCCGGACTATGCAGGATCCTATCCAT ATGACGTTCCAGATTACGCT<u>GAGCTGCGCT</u>CGGAGCTGCCCAGCGTGCCCGGTGC GGCGACAGCAGCAGCGACAGCGACGGGACCACCC-3’, with crRNA sequence underlined and *Dmrta2* ATG initiation codon in bold) into the pronucleus of B6D2F2 zygotes. Viable 2-cell stage embryos were transferred to pseudopregnant CD1 females. Founder mice with the modified *Dmrta2^2xHA^* allele were screened for appropriate integration of the transgenic sequences at the knock-in region as well as for the presence of predicted (http://crispr.dbcls.jp/) potential off-targets followed by PCR followed by Sanger sequencing of the PCR products (Genewiz). Germline transmission of the selected *Dmrta2^2xHA^* founder mouse was then validated by PCR.

The *Dmrta2* null and WT alleles were detected by PCR using primers Fwd: 5’-CGAATCTTTCGGACACTGTAGA-3’; Rev WT: 5’-CCAGAC CCTCAAGCACTCAA-3’; Rev KO: 5’-AGCGCCTCCCCTACCCGGTA-3’. The *Dmrta2^Tg^* allele was detected by PCR using primers Fwd Rosa26 5’-AAACTGGCCCTTGCCATTGG-3’; Fwd eGFP 5’-AACGAGAAGCGCGATCACAT-3’; Rev Rosa26 5’-GTCTTTAATCTACC TCGATGG-3’.

The *Pax6^sey^* mutated allele was detected by high-resolution DNA melting analysis (HRMA) (Reed et al., 2007) using primers 5’-AGGGGGAGAGAACACCAACT-3’ and 5’-CATCTGAGCTTCATCCGAGTC-3’. The *Dmrta2^2XHA^* allele was detected by PCR using primers inside the inserted 2XHA sequence (5’-TATCCCTATGACGTCCCGGAC-3’and 5’-AGCGTAATCTGGAACGTCATAT-3’) and flanking the inserted 2XHA sequence (5’-TTCCAGTTCGTTTCCCCAGCA-3’ and 5’-GTACTTCTCCGCTGCCCTCAA-3’).

### Immunofluorescence and in situ Hybridization

For immunofluorescence (IF), dissected brains of embryos (E12.5) were fixed for 2h at 4 °C in 4% paraformaldehyde (PFA) / phosphate-buffered saline (PBS). Tissues were then rinsed with ice-cold PBS and cryopreserved by overnight (O/N) incubation in 30% sucrose / PBS O/N, and subsequently embedded in gelatin (7.5% gelatin, 15% sucrose / PBS), and sectioned in 14 μm cryostat sections. Sections were then rinsed in PBS, permeabilized, and blocked in a solution of PBS with 0.3% Triton X-100 containing 10% goat serum for at least 2h at RT, and then incubated with primary antibodies diluted in blocking serum O/N, at 4°C. Slides were incubated in secondary antibodies diluted in PBST for 2h at RT. The following primary antibodies were used: rabbit anti-Dmrta2 (1:1000) (De Clercq et al., 2018), rabbit anti-Flag (1:500, Sigma, catalog #F3165); rabbit anti-HA (1:500, Abcam, catalog #ab9110), and mouse anti-Zfp423 (1:500, Santa Cruz, catalog #sc-393904). The following secondary antibodies were used: anti-mouse AlexaFluor-488 (1:400, Invitrogen, catalog #A-11017), anti-rabbit AlexaFluor-488 (1:400, Invitrogen, catalog #A-11008), anti-mouse AlexaFluor-594 (1:400, Invitrogen, catalog #A-11005) and anti-rabbit AlexaFluor-594 (1:400, Invitrogen, catalog #A-11012). Sections were counterstained with DAPI. Images were acquired with a Carl Zeiss LSM 710 confocal microscope using Zen-Black software or Nikon A1R gallium arsenide phosphide inverted Con-focal Microscope and processed using ImageJ and Photoshop software.

*In situ* hybridization (ISH) on whole brains or sections were processed with digoxigenin (Dig)-labeled riboprobes as described (Schaeren-Wiemers and Gerfin-Moser, 1993; Wilkinson & Nieto, 1993). The brains of embryos (E11.5, E12.5, and E15.5) and newborn mice (P7) were fixed in 4% PFA/PBS at 4°C, O/N. For ISH on sections, the brains were infused in 30% sucrose/PBS, O/N, and frozen in gelatin (7.5% gelatin, 15% sucrose/PBS). 20-μm cryostat sections were collected. For ISH on whole-mount, the brains were dehydrated by rinsing 15 minutes (min) in successive baths of MetOH-PTw (25%, 50%, 75%, and 100%) and stored at -20°C. The probes were generated from the following previously described cDNA clones: *Dmrta2* (Saulnier et al., 2013), *Gsx2* (Toresson et al., 2000), *Pax6* (Hamasaki et al., 2004), *Rorβ* (De Clercq et al., 2018), *Wnt3a* (Monuki et al., 2001), *Wnt8b* (Lee et al., 2000), and *Zfp423* (Masserdotti et al., 2010). Images were acquired with an Olympus SZX16 stereomicroscope and an XC50 camera, using the Imaging software CellSens.

### Whole-mount X-gal staining of embryos

Embryos were fixed in 4% PFA for 1h at RT and rinsed in detergent solution (2mM MgCl_2_, 0.01% sodium deoxycholate, 0.02% NP-40 diluted in 0.1 M pH 7.3 phosphate buffer). Embryos were then incubated at 37 °C in a staining solution composed of detergent solution with 5mM K_3_Fe(CN)_6_, 5mM K_4_Fe(CN)_6_, and 1mg/ml of X-gal. Before clearing embryos were progressively dehydrated in increasing concentrations of MetOH. After several washes in 100% MetOH, embryos were incubated in a solution of MetOH: BABB (1:1) before being incubated and stored in 100% BABB. BABB is made of one part benzyl alcohol for two parts of benzyl benzoate.

### RNA isolation and RT-qPCR

The cortex of embryos was dissected in RNAase-free cold 1X PBS and then immediately frozen at -80°C. RNA extraction was performed using the Monarch^®^ Total RNA Miniprep Kit (New England Biolabs, catalog #T2010s) according to the manufacturer’s protocol. cDNA was synthesized starting from 1-2 μg of total RNA using the iScript cDNA Synthesis Kit (Bio-Rad, catalog #1708891). RT-qPCRs were carried out with the Luna^®^ Universal qPCR Master Mix (New England Biolabs, catalog #M3003) using the StepOne Plus Real-Time PCR system (Applied Biosystems). RT-qPCRs primers used were as follows: *Pax6* (5’-AGGGCAATCGGAGGGAGTAA-3’and 5’-CAGGTTGCGAAGAACTCTGTTT-3’); *Emx2* (5’-GTCCACCTTCTACCCCTGG-3’ and 5’-CCACCACGTAATGGTTCTTCTC-3’); *Lhx2* (5’-TGGCAGCATCTACTGCAAAG-3’ and 5’-TGTGCATGTGAAGCAGTTGA-3’); *Wnt3a* (5’-CAGGAACTACGTGGAGATCATGC-3’ and 5’-CATGGACAAAGGCTGACTCC-3’); *Wnt8b* (5’-CAGCTCTGCTGGGGTTATGT-3’ and 5’-CTGCTTGGAAATTGCCTCTC-3’) and *GAPDH* (5’-GTGAAGGTCGGTGTGAACG-3’ and 5’-AGGGGTCGTTGATGGCAACA-3’) used as a reference gene for normalizing gene expression results. Error bars show the standard deviations of at least three independent experiments. To detect by RT-PCR the *Dmrta2* tagged allele in dissected cortices of *Dmrta2^2xHA^*transgenics, primers 5’-TTCCAGTTCGTTTCCCCAGCA-3’ and 5’-GTACTTCTCCGCTGCCCTCAA-3’ were used.

### Plasmids

To generate the pGL4.23-tk-*E60*-luc reporter, a 2350 bp fragment containing the *Pax6 E60* enhancer was amplified by PCR from the *E60* hsp68-LacZ-pA construct (McBride et al., 2011) using primers 5’-TGGCCTAACTGGCCGGTACCTGCAATGCTAGGGATCAAACC -3’ and 5’-TCCTCGAGGCTAGCGAGCTCATCTTGCCGGTCACCTCACTA-3’ and cloned into the NotI and SpeI restriction sites of the pGL4.23-based luciferase reporter plasmid containing a 32bp tk minimal promoter. The mouse *Pax6* expression plasmid was generated by amplifying by PCR the *Pax6* ORF from a pEFX-Pax6 expression vector (obtained from EA. Grove) using primers 5’-AAGAGGACTT<u>GAATTC</u>GCAGAACAGTCACAGCGGAGTG-3’ and 5’-GTTCTAGAGG<u>CTCCAG</u>TTACTGTAATCGAGGCCAGTAC-3’ containing EcoRI and XhoI and cloning it into the corresponding sites of a pCS2-Myc expression vector. The pCDNA3-Myc*-Zfp423* and pCS2-Flag*-mDmrta2* expression plasmids were described previously (Masserdotti et al., 2010). The *mDmrta2 ***α***DM* mutant has been generated by PCR amplifying two overlapping fragments encoding the N-terminal and C-terminal region of *Dmrta2* using primers 5’-CATTCTGCCTGGGGACGTCGGAGC-3’ and 5’-CGTACAGCAATTGCAACTCGCGCGCCTTCTCCGCTGCCCTCAACAGC-3’ for the N-terminal region, and primers 5’-GCGCGCGAGTTGCAATTGCT-3’ and 5’-CCGGGCCCAATGCATTGGCG-3’ for the C-terminal region, and cloning them into the HindIII and NotI sites of the pCS2-Flag vector using the Gibson Assembly® Cloning Kit (New England Biolabs, catalog #E5510S). The *mDmrta2 Mut C-R* mutant was generated similarly by amplifying two fragments encoding the N-terminal and C-terminal region of *Dmrta2* using primers 5’-CATTCTGCCTGGGGACGTCGGAGC-3’ and 5’-CTTCTCCGCTGCCCTCAACAGCAG-3’ for the N-terminal region, and primers 5’-GCGCGCGAGTTGCAATTGCT-3’ and 5’-CCGGGCCCAATGCATTGGCG-3’ for the C-terminal region, and assembling them with a third overlapping fragment generated *in vitro* (IDT, gBlocks®Gene Fragment) encoding the *Dmrta2* DM domain in which an arginine replaces each cysteine.

### Cell culture and dual-luciferase reporter assays

Embryonal carcinoma P19 cells and human embryonic kidney 293T were grown in DMEM medium (Gibco, catalog #61965-059) supplemented with 10% fetal bovine serum (Gibco, catalog #26140079) and 100 U/ml penicillin-streptomycin (Gibco, catalog #15140-122) and maintained in culture flasks at 37 °C under 5% CO_2_. The cells were subcultured when they reached 80% confluency.

For reporter assays, P19 cells were plated in 12-well plates, and after 24h, transfected using the CalPhos^TM^ Mammalian Transfection Kit (Takara, catalog #631312) with the indicated luciferase reporter and expression plasmids, together with a plasmid encoding the *Renilla* luciferase gene. After 48h, cells were washed with 1XPBS, total cellular extracts were prepared, and luciferase activity was measured using the Dual-luciferase Reporter Assay System (Promega, catalog #E1960). Ratios of *luc/Renilla* luminescence were calculated and presented as fold activation for each reporter. The HDAC class I inhibitor Romidepsin (Medchemexpress, catalog #HY-15149) was used at 0.001–0.02μm. It was added 24h after transfection, and 24h before performing the luciferase reporter assays.

### Co-immunoprecipitation and western blot

For co-immunoprecipitation (Co-IP), cells were lysed in IPH buffer (20mM Tris-HCl pH7.5, 150mM NaCl, 2mM EDTA, 1% NP-40) supplemented with protease inhibitor cocktail (Roche, cOmplete^TM^ EDTA-free Protease Inhibitor Cocktail, catalog # 05892791001) and incubated for 15 min, at 4°C. Cell lysates were then centrifuged twice at 13,000 rpm for 30 min, at 4 °C to remove debris. Part of the lysate (2.5%) was kept as a positive input control. Co-IPs were performed by incubating cell lysates (1 mg) with 5 μl of the indicated antibodies at 4 °C with rotation, O/N. 20 μl of magnetic beads G (Cell Signaling, catalog # 9006s) was then added for 4h, at 4 °C. After four washes with IPH buffer, bound protein complexes were eluted by incubating beads in Laemmli sample buffer for 10 min, at 90°C. Immunoprecipitated protein complexes were subjected to western blot analysis as described below.

For western blot analysis, protein extracts from dissected cortices were prepared in RIPA buffer (50mM Tris-HCl pH8, 150mM NaCl, 1% NP-40, 0.5% sodium deoxycholate, 0.1% SDS,) and those from cultured cells were prepared in IPH buffer supplemented with protease inhibitor cocktail. Homogenates were quantified by using the Bradford Assay (OD 595nm) and centrifuged for 15 min, at 10,000 g, at 4°C, and the supernatants were collected. Equal quantities of protein (60 μg/lane) were separated by 8% to 10% SDS-PAGE in a chamber filled with running buffer (0.1% SDS, 25 mM Tris base, 190 mM glycine; adjust pH to 8.3) and placed into a transfer cassette with transfer buffer (25mM Tris-base, 190 mM glycine, 20% MetOH; adjust pH to 8.3), and the proteins were transferred onto a polyvinylidene difluoride (PVDF, Cytiva, catalog # 10600029) membrane. Then the membrane was blocked with Tris-buffered saline (TBS) with 5% non-fat milk for 1h, at RT and incubated at 4°C, O/N, with the following primary antibodies: rabbit anti-Flag (1:1000); rabbit anti-HA (1:1000); rabbit anti-H3 (1:50,000, Millipore, catalog #07-690,); mouse anti-Myc (1:1000, Sigma, catalog #M4439); mouse anti-Zfp423 (1:1000) and the NurD complex antibody kit (Cell Signaling, catalog #8349T). Following rinsing three times with TBS/Tween-20, the membranes were incubated with the following secondary antibodies: anti-mouse IgG HRP (1:5,000, Jackson ImmunoResearch, catalog #115-035-062) and anti-rabbit IgG HRP (1:3,000, Cell Signaling, catalog #7074) for 1h, at RT. Protein levels were normalized to H3, and all proteins were detected using the NovexTM ECL Chemiluminescent Substrate Reagent Kit (Invitrogen, catalog #WP20005).

### Electrophoretic Mobility Shift Assays

EMSAs were performed using nuclear extracts prepared from HEK293T cells transfected with expression vectors of either DMRTA2 or DMRTA2^R116P^ as described (Dignam et al., 1983). Total protein concentrations were determined by Bradford assays (Bio-Rad). DMRTA2 WT and DMRTA2^R116P^ proteins were also produced using the Transcription and Translation (TnT) system (Promega). EMSAs were performed as described previously (Van Lint et al., 1994). Briefly, 2μl of TNT preparation or 1μg nuclear extracts from HEK293T cells either non-transfected or transfected with DMRTA2 WT or DMRTA2 mutated were first incubated for 10 min in the absence of a probe in a reaction mixture containing 10 μg of DNase-free BSA (Bioké), 2 μg of poly(dI–dC) (Sigma) as non-specific competitor DNA, 50 μM ZnCl_2_, 0.25 mM DTT, 20 mM HEPES (pH 7.3), 60 mM KCl, 1 mM MgCl_2_, 0.1 mM EDTA and 10% (v/v) glycerol. 30 000 c.p.m. of probe (10–40 fmol) was then added to the mixture that was incubated for 20 min. Samples were subjected to electrophoresis at room temperature on 6% polyacrylamide gels at 120V for 2–3 h in 1× TGE buffer (25 mM Tris-acetate (pH 8.3), 190 mM glycine, and 1 mM EDTA). Gels were dried and autoradiographed for 48–72h at −80°C. The probe used is oligonucleotide 5RE described in Murphy et al., 2007.

### Rapid immunoprecipitation mass spectrometry of endogenous protein

Rapid immunoprecipitation mass spectrometry of endogenous protein (RIME) experiments were carried out as previously described (Mohammed et al., 2016) with minor modifications. Briefly, E12.5 cortices were dissected on ice in RNAase-free ice-cold 1XPBS and fixed for 15 min in 1% formaldehyde (Thermos Scientific, catalog #28906) in 1XPBS at RT. The cross-linking was quenched by adding glycine to a final concentration of 125mM and incubating samples for 8 min. Subsequently, the fixed tissues were pelleted and washed in ice-cold PBS. Nuclear extraction was done using ice-cold LB1 buffer (50mM HEPES-KOH pH7.5, 140mM NaCl, 1mM EDTA, 10% glycerol, 0.5% NP-40 (vol/vol) and 0.25% Triton X-100(vol/vol)), incubating cells on rotation for 10 min at 4°C. The suspension was then centrifuged at 2000g for 5 min at 4°C and the obtained nuclear pellet was washed in ice-cold LB2 buffer (10 mM Tris-HCl pH 8.0, 200 mM NaCl, 1 mM EDTA, 0.5 mM EGTA). Pelleted nuclei were subsequently resuspended in LB3 buffer (10 mM Tris-HCl pH 8.0, 100 mM NaCl, 1mM EDTA, 0.5 mM EGTA, 0.1% sodium deoxycholate (wt/vol) and 0.5% N-lauroylsarcosine (wt/vol)) and, after an incubation of 15 min, were subjected to 3 cycles of sonication (30sec ON/30sec OFF for 10 min, at 200W, high setting, Bioruptor, Diagenode). Triton X-100 was added to the sonicated lysate to a final concentration of 1%. The obtained chromatin was then cleared by centrifugation at 20,000g for 10 min, at 4 °C, and the supernatant was collected. All lysis buffers were supplemented with protease inhibitors before use.

Antibody-bound beads were generated by incubating 100 μl Protein G magnetic beads (Cell Signaling, catalog #cat 9906) with 5 μl rabbit anti-HA (Abcam, catalog #ab9110,) or rabbit anti-IgG (Merck, catalog #12-370) in PBS/BSA and incubated in rotation at 4 °C, O/N. The next day, the antibody-bound beads were washed three times in PBS/BSA to remove any unbound antibodies. For immunoprecipitation, 2 mg of protein lysate was incubated O/N, with 100 µl antibody-bound beads at 4 °C. The next day, the bead-bound complexes were washed nine times in RIPA buffer (50 mM HEPES pH 7.6, 1 mM EDTA, 0.7% sodium deoxycholate, 1% NP-40, and 0.5M LiCl) at 4 °C and then washed two times with a cold 100 mM ammonium hydrogen carbonate (AMBIC) solution. Finally, the beads were transferred to new tubes, snap-frozen at -80 °C, and stored at this temperature until mass spectrometry analysis.

### Mass spectrometry

The samples were treated using Filter-aided sample preparation (FASP). To first wash the filter, 100 µl of 1 % formic acid was placed in each Millipore Microcon 30 MRCFOR030 Ultracel PL-30 and samples were centrifuged for 15 min, at 14500 rpm. For protein adjustment, 40 µg of protein in 150 µl of 8M urea buffer (urea 8 M in 0.1 M Tris, pH 8,5) were placed individually in a column and centrifuged for 15 min, at 14500 rpm. The filtrate was discarded, and the columns were washed three times with 200 µl of urea buffer followed by centrifugation for 15 min, at 14500 rpm. For the reduction step, 100 µl of dithiothreitol (DTT) was added and mixed for 1 min, at 400 rpm with a thermomixer, incubated for 15 min, at 24 °C, and centrifuged for 15 min, at 14500 rpm. The filtrate was discarded and 100 µl of urea buffer was added before another centrifugation for 15 min, at 14500 rpm. An alkylation step was performed by adding 100 µl of iodoacetamide (IAA), in urea buffer in the column, mixing for 1 min, at 400 rpm, incubating for 20 min in the dark, and centrifuging for 10 min, at 14500 rpm. To remove the excess IAA, 100 µl of urea buffer was added and the samples were centrifuged for 15 min, at 14500 rpm. To quench the remaining IAA, 100 µl of DTT was added to the column, mixed for 1 min, at 400 rpm, incubated for 15 min, at 24 °C, and centrifuged for 10 min, at 14500 rpm. Excess DTT was removed by adding 100 µl of urea buffer and centrifuging for 15 min, at 14500 rpm. The filtrate was discarded, and the column was washed three times with 100 µl of sodium bicarbonate buffer 50 mM (ABC) followed by centrifugation for 10 min, at 14500 rpm. The last 100 µl were kept at the bottom of the column to avoid any evaporation in the column. For digestion, 80 µl of mass spectrometry grade trypsin (1/50 in ABC buffer) was added to the column, mixed for 1 min, at 400 rpm, and incubated at 24°C, O/N in a water-saturated environment. The Microcon columns were placed on a LoBind tube of 1.5 ml and centrifuged for 10 min, at 14500 rpm. 40 µl of ABC buffer was added to the column before centrifugation for 10 min, at 14500 rpm. 10% Trifluoroacetic acid (TFA) was added to the content of the LoBind Tube to obtain 0.2 % TFA. The samples were dried in a SpeedVac up to 20 µl and transferred for injection.

The samples were analyzed using nano-LC-ESI-MS/MS (timsTOF Pro, Bruker, Billerica, MA, USA) coupled with a UHPLC nanoElute (Bruker). Liquid chromatography was separated at 50°C and at a flow rate of 200 nl/min by nanoUHPLC (nanoElute, Bruker) on a C18 column (25 cm x 75 μm ID) with integrated CaptiveSpray insert (Aurora, ionopticks, Melbourne). Mobile phases A and B were water with 0.1% formic acid (v/v) and ACN with formic acid 0.1% (v/v), respectively. Samples were loaded directly on the analytical column at a constant pressure of 800 bar. The digest (1 µl) was injected, and the organic content of the mobile phase was increased linearly from 2% to 15 % B within 40 min, followed by an increase to 25% B within 15 min, and further to 37 % B in 10 min, followed by a washing step at to 95% B in 5 min. Data acquisition on the timsTOF Pro was performed using Hystar 6.1 and tims Control 2.0. The tims TOF Pro data were acquired using 160 ms TIMS accumulation time, mobility coefficients (1/K0) range from 0.75 to 1.42 Vs/cm². Mass-spectrometry analysis was carried out using the parallel accumulation-serial fragmentation (PASEF) (Meier et al., 2018) acquisition method. One MS spectra followed by PASEF MSMS spectra per total cycle of 1.16s. All MS/MS samples were analyzed using Mascot (Matrix Science, London, UK; version 2.8.1). Scaffold (version Scaffold_5.0.0, Proteome Software Inc., Portland, OR) was used to validate MS/MS-based peptide and protein identifications.

### Human subjects and clinical evaluation

As a result from a GeneMatcher match (Sobreira et al., 2015), we studied a Pakistani consanguineous family with three affected individuals, V-3, V-5, and V-7 (Fig. 6Ac). The clinical evaluation of the probands was performed at the (Authors’ University Hospital). Detailed clinical evaluation was carried out in two affected persons, V-3, and V-5 (**Extended data Table 6.1**), however, the affected person V-7 showed similar clinical features. This study was conducted with ethical approval from the (Authors’ University). We obtained written informed consent, and blood samples were taken from all the available persons (III-7, IV-1, IV-5, IV-7, V-3, V-5, V-7) in the family. Genomic DNA was extracted as described previously (Sambrook et al., 2006). The MRI of V-3 (6 years) was performed, which revealed microcephaly, a simplified gyral pattern along with agenesis of the corpus callosum and colpocephaly. Electroencephalography (EEG) was performed on the affected person, V-5 (5 years).

### Whole-exome sequencing

We performed whole-exome sequencing (WES) using DNA samples of two affected individuals, V-3, and V-5 (**Fig. 6 Ac**). WES and further data analysis to find the exact causative variant has been performed as described (Yousaf et al., 2022, 2023). The combined analysis of WES data of V-3 and V-5 revealed three shared homozygous variants; *C1orf109*: NM_001303030.2: c.373C>A:p. (Leu125Ile), *DMRTA2*: NM_032110.3: c.347G>C:p.(Arg116Pro), *TMEM161A*: NM_001256766.3: c.745G>A:p.(Ala249Thr) between both the WES-analyzed persons. We further excluded two variants located in *C1orf109* and *TMEM161A* genes, as the variant in *C1orf109* was found 11x in homozygous state in the gnomAd database (v4.1.0), and the *TMEM161A* variant did not segregate with the phenotype. *DMRTA2*: p. (Arg116Pro) is the only homozygous variant that segregated (tested by Sanger sequencing) with the phenotype in this consanguineous family (**Fig. 6**). The variant p. (Arg116Pro) has a Combined Annotation Dependent Depletion CADD score (v1.6) of 31 (Kircher et al., 2014), and is predicted as “deleterious” by multiple in silico-prediction programs and classified as a variant of uncertain significance, according to the ACMG classification criteria PM2, PP1, PP3, and PP4 (Richards et al., 2015).

### Statistical analysis

Statistical significance was determined by the Student’s *t-*test, with a threshold for significance set to p < 0.05. All results are plotted as the mean ± S.D., as indicated in figure legends.

## RESULTS

### *Dmrta2* cooperates with *Pax6* in defining telencephalic dorsoventral compartments and acts through its repression to control cortical patterning

To test the hypothesis that *Dmrta2* acts through *Pax6* to control cortex development, we generated double mutants by intercrossing *Dmrta2^+/-^* mice with *Pax6^sey/+^* mice which have a point mutation in the *Pax6* gene creating a null allele (Hill et al., 1991). Embryos were collected at three embryonic time points (E11.5, E15.5, and E18.5) to measure the surface area of their hemispheres. We found that the cerebral hemispheres of *Pax6^sey/sey^* embryos and *Dmrta2^-/-^* embryos were smaller compared to embryos at E18.5, with the reduction in *Dmrta2*^-/-^ being more severe (-60.1% ± 2.5% in *Dmrta2^-/-^* compared to -16.9% ±7.2% in *Pax6^sey/sey^*) and detected at an earlier stage in *Dmrta2^-/-^*embryos than in *Pax6^sey/sey^* embryos (**Extended data Fig. 1-1**). In the *Dmrta2^-/-^; Pax6^sey/sey^* embryos, the cerebral hemispheres were virtually absent. Notably, the extent of the reduction in *Dmrta2^-/-^* embryos was less severe in the absence of one allele of *Pax6* (-31.6% ± 6.5% in *Dmrta2^-/-^; Pax6^sey/+^* compared to -60.1% ± 2.5% in *Dmrta2*^-/-^) (**Fig. 1**). This rescue was already visible at earlier stages: at E11.5 (-38.6 % ± 5,6% in *Dmrta2^-/-^; Pax6^sey/+^* compared to -48.2% ± 6.4% in *Dmrta2^-/-^*) and E15.5 (-34.5 % ± 4.2% in *Dmrta2^-/-^; Pax6^sey/+^* compared to -48.9% ± 7.7% in *Dmrta2^-/-^*) (**Extended data Fig. 1-1**). Analysis by *in situ* hybridization of the expression of the subpallial marker *Gsx2* revealed that its expression in the single KO embryos remained confined to the subpallium or only very slightly crossed the pallium-subpallium boundary (PSB) while it extended into the abortive cortical primordium of the double KO embryos (**Fig. 2**). These data indicated that *Dmrta2* and *Pax6* cooperate in maintaining cortical identity in dorsal telencephalic progenitors.

**Figure. 1.**
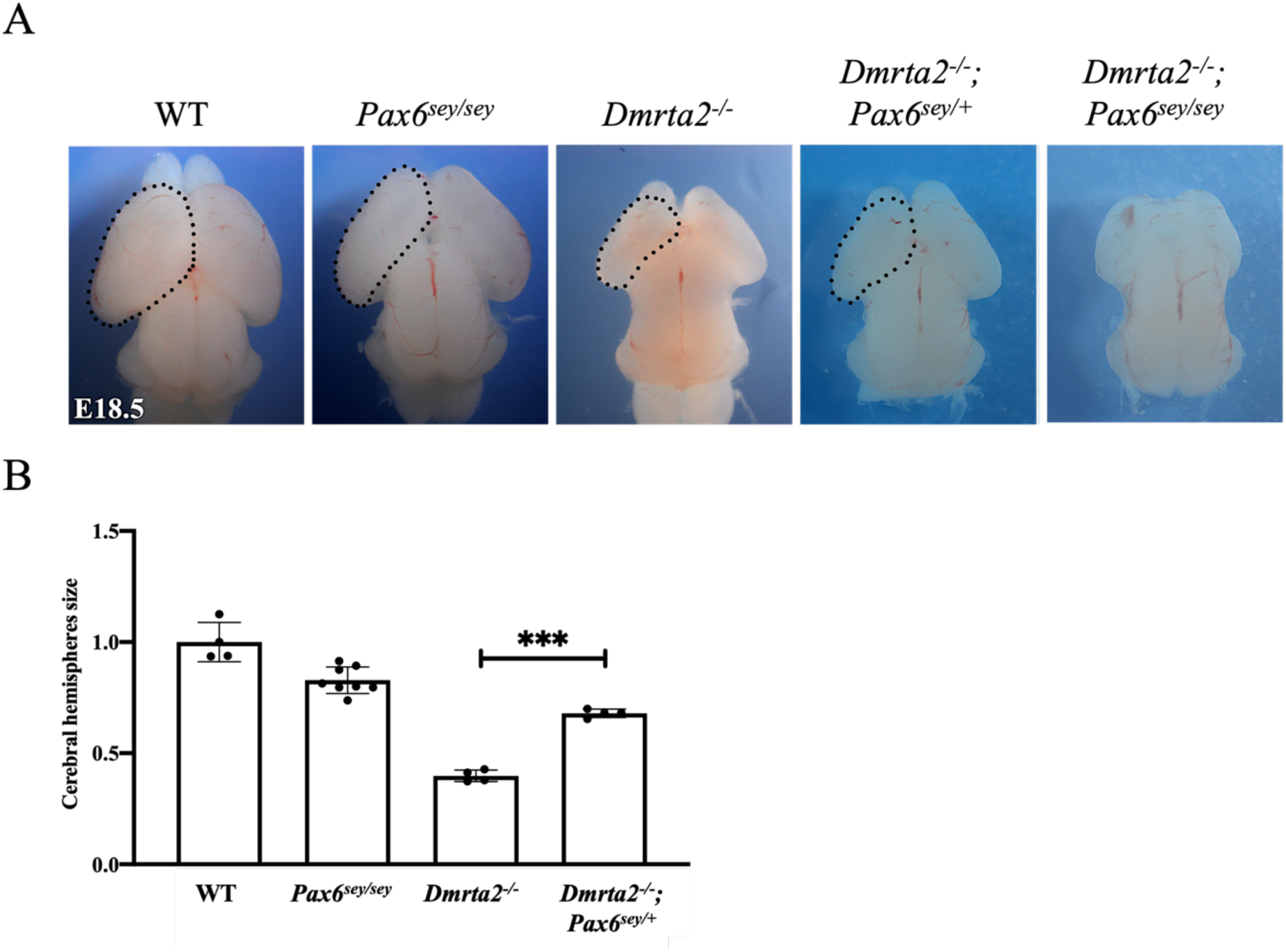
Reduction of *Pax6* partially rescues telencephalic vesicle growth in *Dmrta2^-/-^* embryos. **(A)** Dorsal views of the brain of E18.5 embryos of the indicated genotypes. **(B)** Graphs representing the measured surface area of the cerebral hemispheres of *Pax6^sey/sey^*, *Dmrta2^-/-^* and *Dmrta2^-/-^; Pax6^sey/+^* compared to WT set to 1. ***P < 0.001.

**Figure 2.**
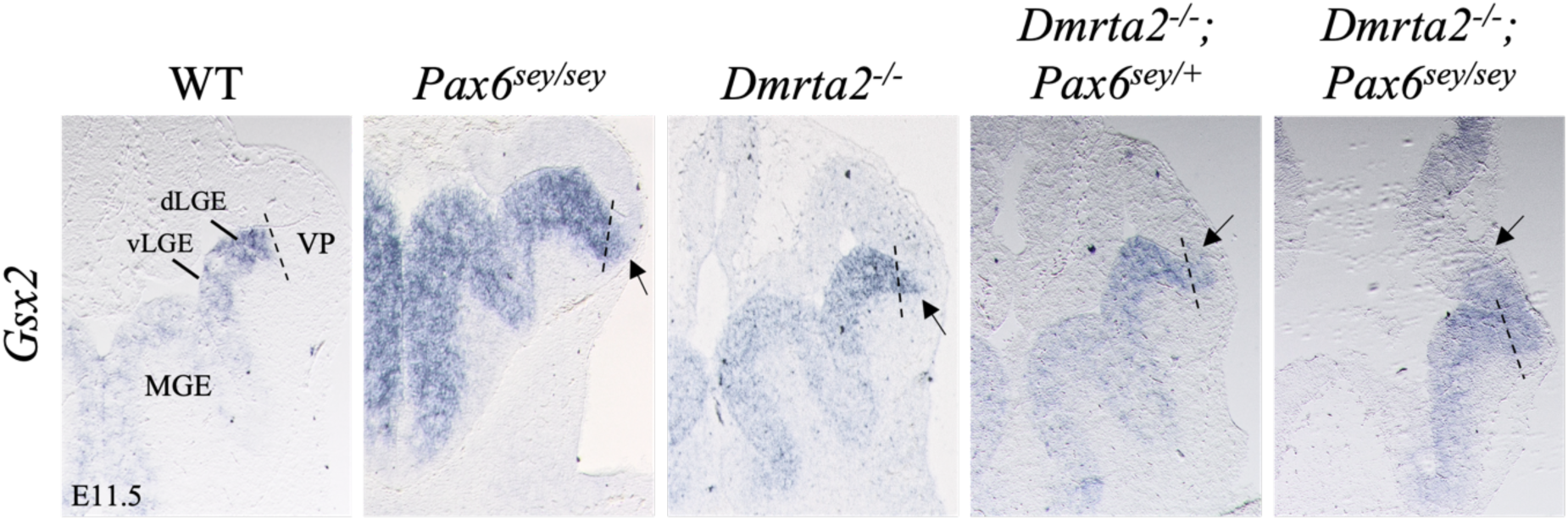
The ventral determinant *Gsx2* expands dorsally in the reduced cortical primordium of *Dmrta2^-/-^; Pax6^Sey/Sey^* embryos. Coronal sections through the brain of E11.5 embryos of the indicated genotypes processed by ISH showing *Gsx2* expression in the telencephalon. Note that *Gsx2* only slightly crosses the pallium-subpallium boundary (PSB, indicated by dashed lines) in *Pax6^sey/sey^* and *Dmrta2^-/-^* embryos, slightly expands into the lateral dorsal telencephalon of *Dmrta2^-/-^; Pax6^sey/+^*embryos and more robustly in *Dmrta2^-/-^; Pax6^sey/sey^* embryos. Arrows point to the limit of the expansion of *Gsx2* expression within the dorsal telencephalon.

*Pax6* suppresses medial cortical fate in the cortical neuroepithelium (Godbole et al., 2017). Therefore, and because *Pax6* is excluded from the hem (Konno et al., 2002; Saulnier et al., 2013; De Clercq et al., 2018), we hypothesized that the medial expansion of *Pax6* observed upon the loss of *Dmrta2* could be responsible for the suppression of hem formation. To test this hypothesis, we examined by *in situ* hybridization the expression of *Wnt3a* which identifies the cortical hem, and *Wnt8b,* which marks the dorsomedial telencephalic primordium and eminentia thalami, both being upregulated in the absence of *Pax6* (Godbole et al., 2017), in the cortex of E11.5 homozygous and heterozygous *Dmrta2* mutants with the loss of one *Pax6* allele (**Fig. 3A**). As previously reported (Saulnier et al., 2013), in the medial telencephalon of *Dmrta2^-/-^* embryos, *Wnt3a* was strongly reduced. *Wnt3a* was much less affected in *Dmrta2^-/-^; Pax6^sey/+^* embryos than in *Dmrta2^-/-^* embryos and some expression could be also detected medially in the reduced cortex of *Dmrta2^-/-^; Pax6^sey/sey^* embryos. A slight reduction in the intensity of *Wnt3a* expression was also observed in *Dmrta2^+/-^* embryos, while an expansion was detected in *Dmrta2^+/-^; Pax6^sey/+^* embryos, even more pronounced in *Dmrta2^+/-^; Pax6^sey/sey^* embryos. In *Dmrta2^-/-^* embryos, as reported previously, *Wnt8b* was also downregulated in the telencephalon but not in the eminentia thalami (Saulnier et al., 2013). In *Dmrta2^-/-^; Pax6^sey/+^*embryos, *Wnt8b* expression appeared to extend slightly more dorsally in the reduced medial telencephalon than in *Dmrta2^-/-^* embryos; and appeared, in contrast, more restricted in the abortive cortex of *Dmrta2^-/-^; Pax6^sey/sey^* embryos. In *Dmrta2^+/-^* embryos, *Wnt8b* appeared unchanged compared to controls. However, its expression expanded dorsally in *Dmrta2^+/-^; Pax6^sey/+^* embryos and this expansion was even more severe in *Dmrta2^+/-^; Pax6^sey/sey^* embryos. RT–qPCR on RNA extracted from telencephalic tissue isolated from E11.5 WT and mutants confirmed the drastic downregulation of *Wnt3a and Wnt8b* in *Dmrta2^-/-^* embryos and its rescue upon the loss of one *Pax6* allele (**Fig. 3B**). Thus, patterning of the cerebral cortex and cortical hem formation is regulated by *Dmrta2* through repression of *Pax6* which restricts medial cortical fate to the medial telencephalon.

**Figure 3.**
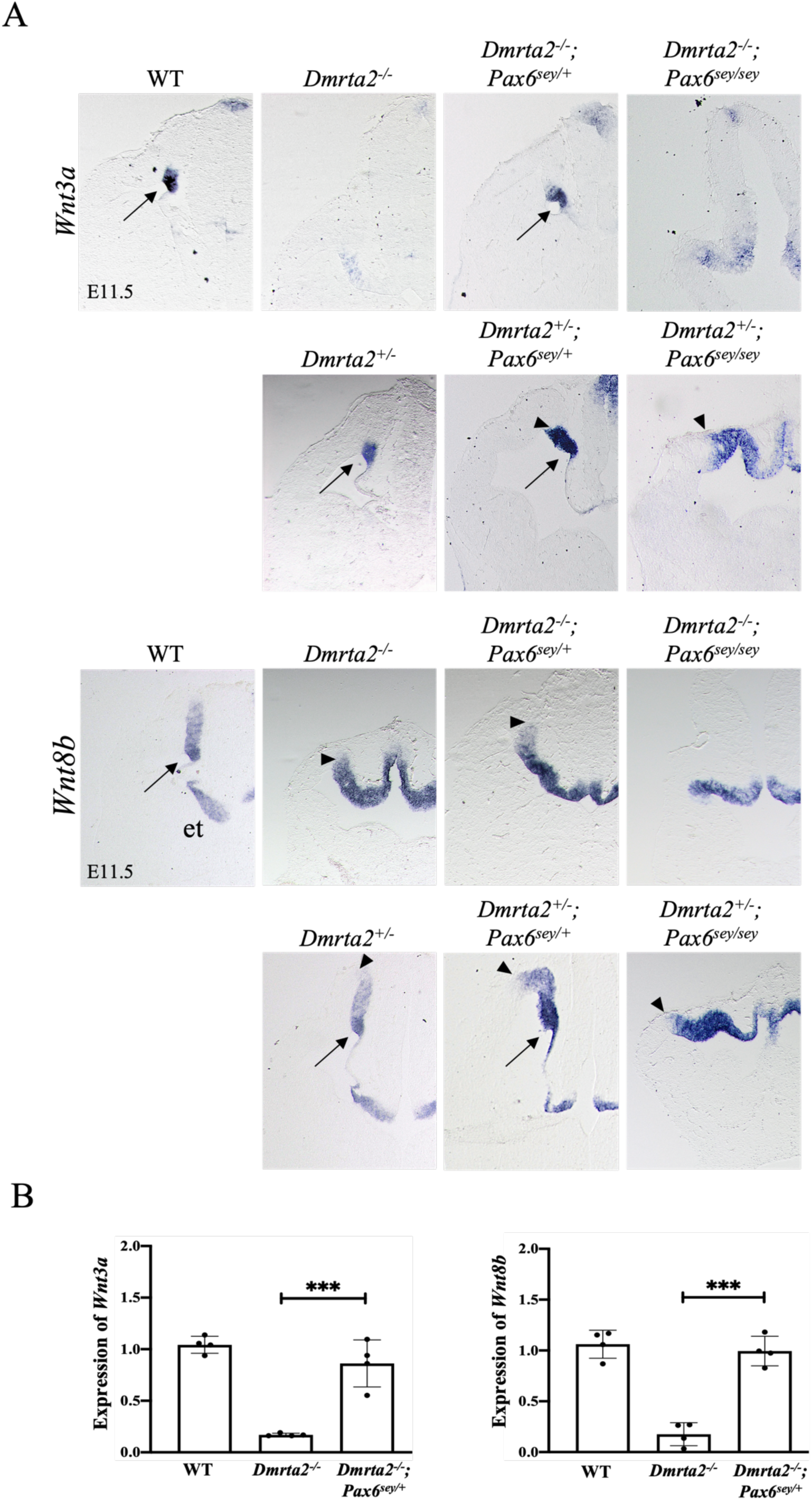
Reduction of *Pax6* partially rescues medial cortical fate in *Dmrta2* homo- and heterozygous mutant embryos. **(A)** Coronal sections through the brain of E11.5 embryos of the indicated genotype processed by ISH for the expression of *Wnt3a* marking the hem and *Wnt8b* marking the dorsomedial telencephalic primordium. Arrows point to the hem. Arrowheads indicate the limit of the expansion of the expression of these markers into the dorsal pallium. Et: eminentia thalami. **(B)** Quantitative RT–qPCR analysis of *Wnt3a* and *Wnt8b* in the cortex of *Dmrta2^-/-^*, *Dmrta2^-/-^; Pax6^sey/+^*and WT control embryos. Results are normalized to the level of the expression detected in the cortex of WT embryos. ***P < 0.001.

*Pax6* promotes rostral area identity in the developing neocortex (Bishop et al., 2000, 2002; O’Leary and Sahara, 2008). *Pax6* is reduced in the cortex of transgenic mice conditionally overexpressing *Dmrta2* that have expanded V1 area (De Clercq et al., 2018). It could therefore be that it is the decreased level of *Pax6* that caudalizes the neocortex of mice expressing in excess *Dmrta2*. Therefore, we generated transgenic mice conditionally overexpressing both *Dmrta2* and *Pax6* by crossing *Emx1^Cre^; Dmrta2^Tg/+^* (De Clercq et al., 2018) with *Emx1^Cre^*; *Pax6^Tg/+^* mice (Berger et al., 2007). We examined area formation at P7 in the cortex of heterozygous double transgenics (*Emx1^Cre^; Dmrta2^Tg/+^; Pax6^Tg/+^*) and single transgenic mice (*Emx1^Cre^; Dmrta2^Tg/+^* and *Emx1^Cre^; Pax6^Tg/+^*), as homozygous ones exhibit early lethality, possibly due to sustained expression of *Dmrta2* and *Pax6* in post-mitotic cortical cells. This analysis was carried out by whole-mount *in situ* hybridization monitoring *Rorβ* expression, which demarcates the S1, A1, and V1 area, measuring the size of the V1 area relative to the total hemisphere size (**Fig. 4**). We found that the V1 area that are expanded in *Dmrta2* overexpressing mice are reduced when *Pax6* is also overexpressed, as observed in *Pax6* overexpressing mice. These findings support the hypothesis that *Dmrta2* controls cortical arealization by regulating the level of *Pax6* expression.

**Figure 4.**
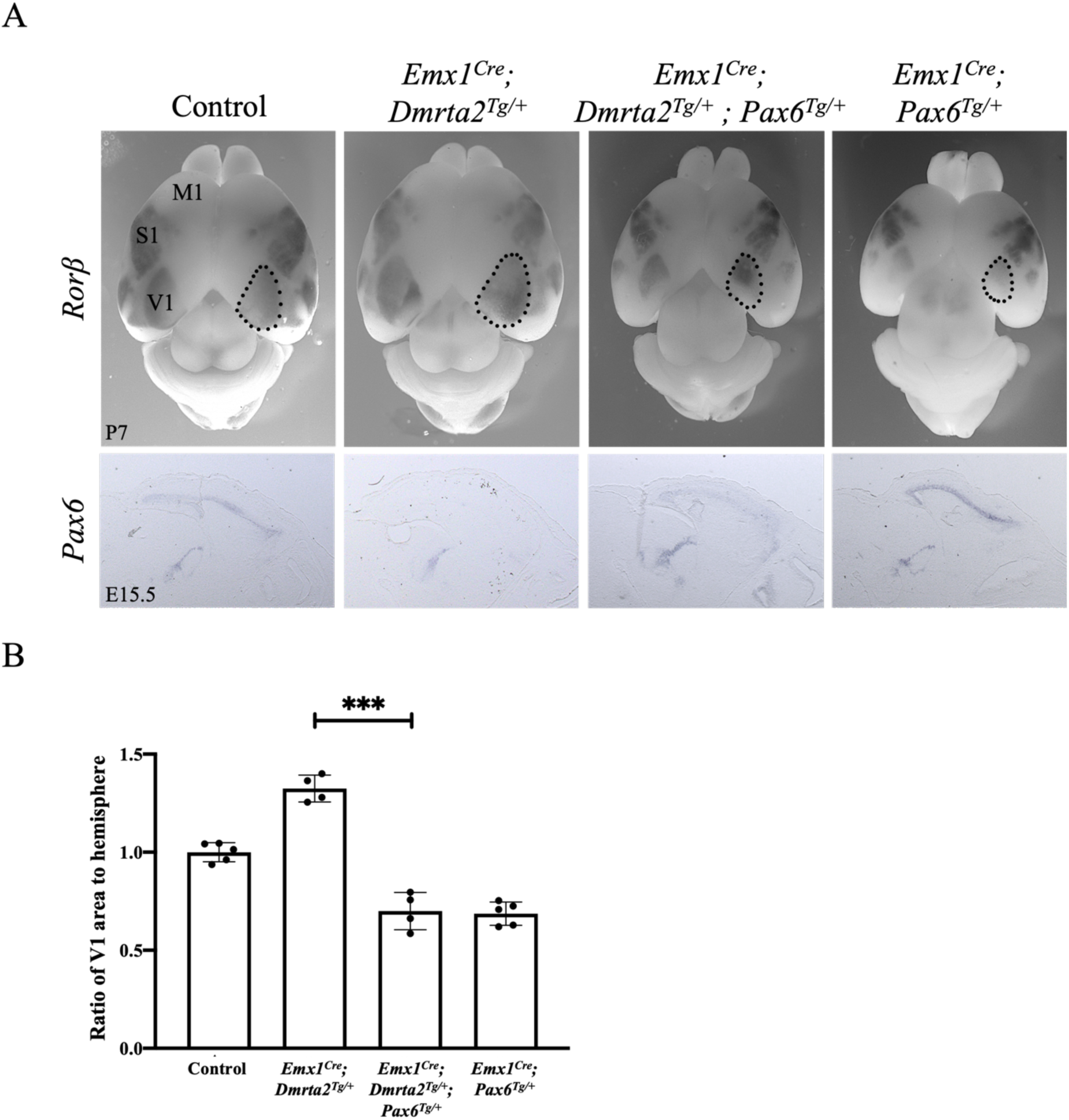
*Pax6* overexpression rescues the expansion of the V1 area observed in *Dmrta2* overexpressing transgenics. **(A)** Dorsal views of the brain of P7 neonates of the indicated genotype processed by whole-mount ISH for *Rorβ* expression. *Dmrta2^Tg/+^; Pax6^Tg/+^* were used as controls. Sagittal sections of the brain of E15.5 embryos of the corresponding genotype with *Pax6* expression detected by ISH are shown below. M1: primary motor area; S1: primary somatosensory area; V1:primary visual areas. **(B)** Histograms show that as a ratio of surface area to total hemisphere size, V1 expansion is significantly rescued by the reduction of *Pax6*. ***P < 0.001.

### Dmrta2 can act as a DNA-binding repressor on the *Pax6 E60* enhancer and a point mutation in its DM domain causes microcephaly in human

*Pax6* has been shown to be deregulated by the loss or gain-of-function of *Dmrta2* in the developing cortex (Konno et al., 2012; Saulnier et al., 2013). The deregulation of *Pax6* is rapid as it is already observed at E12.5 in *Nestin^Cre^; Dmrta2^fl/fl^* mice (De Clercq et al., 2018). It can be seen already at E9.5 in *Dmrta2* null mutants before the disruption of Wnt signaling that can be detected from E10.5 in Wnt reporter *BAT-gal* mice (Maretto et al., 2003) (**Extended data Fig. 5-1**) suggesting *Pax6* is a direct Dmrta2 target. In accordance, ChIP-seq experiments have revealed that Dmrta2 binds to the *Pax6 E60* enhancer, an ultra-conserved cis-regulatory region located 25kb downstream of *Pax6* in the large final intron of the adjacent *Elp4* gene, that contains several potential Dmrta2 binding sites and drives complex expression of *Pax6* within the nervous system, including in the telencephalon (McBride et al., 2011; Konno et al., 2019). To address the possibility that Dmrta2 affects *Pax6 E60* enhancer activity and investigate its mechanism of action, we performed transfection experiments in P19 teratocarcinoma cells with a luciferase reporter construct driven by a minimal tk promoter linked to the *Pax6 E60* enhancer. As Pax6 autoregulates its expression (Aota et al., 2003), we transfected either naive or *Pax6* overexpressing cells with this reporter, in the absence or presence of *Dmrta2* expression vectors. We found that while Pax6 overexpression had no significant effect on the control enhancer-less tk luciferase reporter, it increased the activity of the *E60* enhancer. In the presence of Pax6, we found that Dmrta2 represses the activity of the *E60* enhancer in a dose-dependent manner (**Fig. 5A**). Such repression was not observed with cotransfection of a construct encoding a *Dmrta2* mutant (with the cysteine residues of the zinc finger motif replaced by arginine residues, designated *Dmrta2 mut C-R*) that without affecting its subcellular localization abolishes its ability to bind DNA (**Fig. 5B and Extended data Fig. 5-2**). These findings indicated that Dmrta2 functions as a DNA-binding transcription repressor and suggest that it may attenuate *Pax6* by blocking its autoregulation.

**Figure 5.**
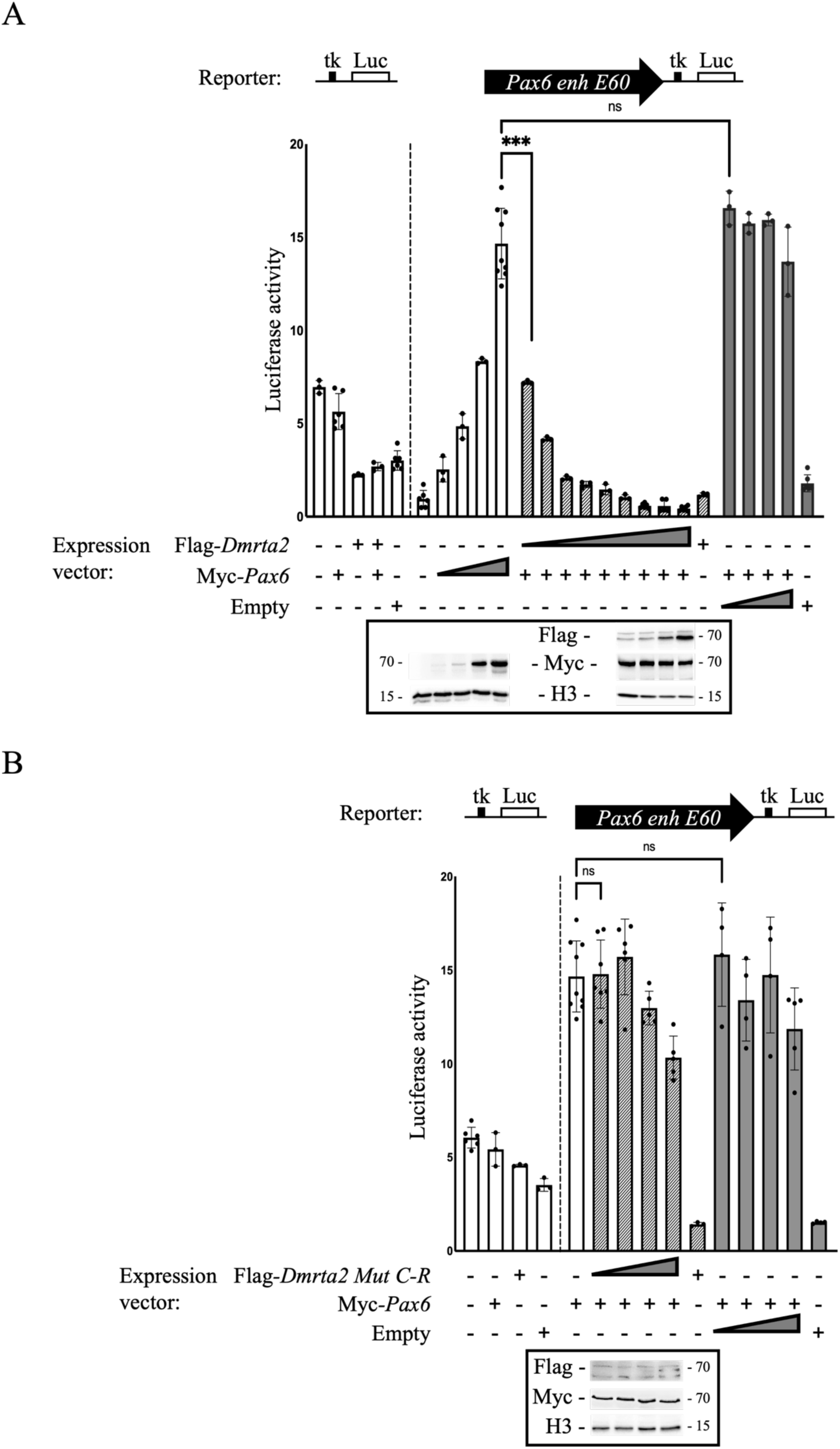
*Dmrta2* inhibits the activation by *Pax6* of the *Pax6 E60* enhancer. **(A)** Reporter assays in P19 cells transfected with a *Pax6 E60* tk-luciferase reporter vector, or an ‘empty’ tk-luciferase reporter vector as indicated, together with a Myc*-Pax6* expression vector and/or a Flag*-Dmrta2* expression vector at increasing doses, as indicated. Note that the *Pax6* dose-dependent transactivation of the *E60* tk-luciferase reporter vector is attenuated by *Dmrta2*. Western blots showing the expression levels of Myc*-Pax6* and Flag*-Dmrta2* are shown below. **(B)** Reporter assays in P19 cells transfected with a *Pax6* E60 tk-luciferase reporter vector or an ‘empty’ tk-luciferase reporter vector as indicated, with a Myc*-Pax6* expression vector and/or a Flag*-Dmrta2* point mutant with the cysteine residues of the zinc finger motif replaced by arginine residues (Flag*-Dmrta2 Mut C-R*) at increasing doses, as indicated. Western blots showing the expression levels of Myc*-Pax6,* and Flag*-Dmrta2 Mut C* are shown below. Note that in contrast to the WT protein, the Flag*-Dmrta2 Mut C-*R is unable to repress *Pax6* transactivation of the *E60* reporter construct.

Besides, by exome sequencing, we identified a homozygous novel single base pair deletion (c.1197delT) predicted to result in a frameshift variant p. (Pro400Leufs*33) in *DMRTA2* in a consanguineous family with three siblings affected by a severe prenatal neurodevelopmental disorder characterized by frontoparietal pachygyria, agenesis of the corpus callosum and progressive severe microcephaly (Urquhart et al., 2016). This is, to our knowledge, the only case report suggesting that loss of DMRTA2 causes cortical malformation in humans. Independent reporting of biallelic variants in *DMRTA2* in individuals with the same or similar phenotypes is required to attain the level of evidence required for a definite gene-disease relationship. Here, we identified a homozygous missense variant in the DM domain of *DMRTA2*:NM_032110.3:c.347G>C:p.(Arg116Pro) that segregated with cortical malformations in three branches of a consanguineous family, and arginine at position 116 is extremely conserved down to *C.elegans*, suggesting that the *de novo* DMRTA2^R116P^ mutation is the most likely cause of the phenotype (**Fig. 6A**). Patients presented global developmental delay, dysarthria, muscle atrophy, aggressive behavior, peripheral neuropathy, and were unable to stand and walk. MRI analysis in one affected person (V-3) revealed microcephaly, a complete absence of the corpus callosum, ventriculomegaly in the posterior part of the body, and colpocephaly (**Fig. 6B**). Electroencephalogram (EEG) was performed in V-5, which revealed abnormalities indicating a mild encephalopathy. Full details of the clinical and genetic characterization of the V-3 and V-5 patients are provided in **Extended data Table 6-1.**

**Figure 6.**
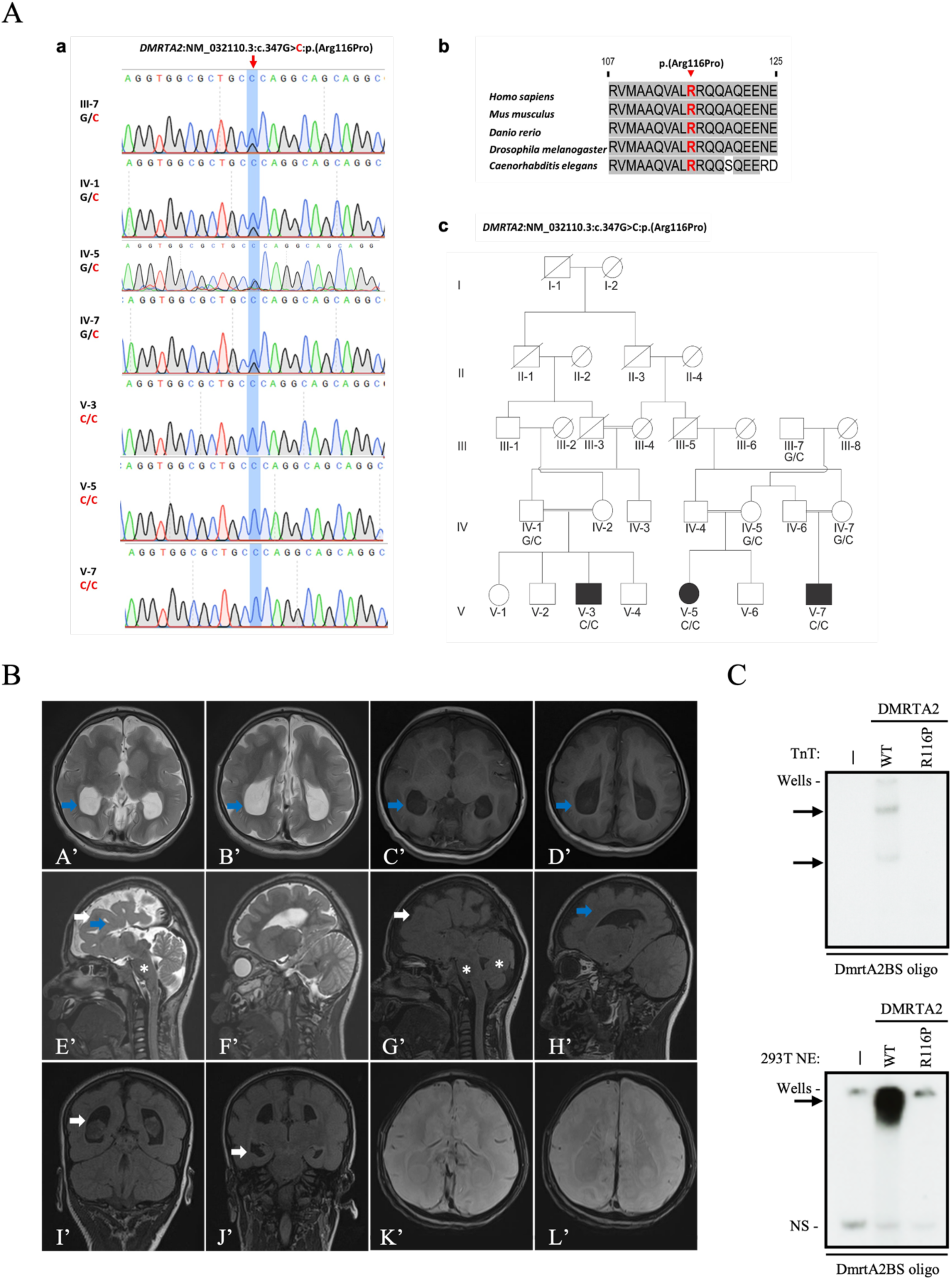
The homozygous missense variant in *DMRTA2* causes microcephaly in human. **(A)** Sanger sequencing chromatograms **(a)** showing *de novo* G to C heterozygous and homozygous changes in sequence causing p.Arg116Pro coding change in *DMRTA2* identified in individuals of a consanguineous family. An alignment of the sequence of the central portion of the major groove recognition helix of human, murine fish, *Drosophil*a, and *Caenorhabditis elegans Dmrta2* is shown **(b)**, together with the pedigree **(c)** of the Pakistan family in which the mutation has been identified. Symbols marked by a slash indicate that the subject is deceased. Males are indicated by squares; females are indicated by circles. Blackened symbols represented individuals who were identified as patients according to clinical examination. **(B)** Magnetic resonance (MRI) images of the brain phenotype. First Row: Axial T2 weighted (A’, B’) and T1 weighted (C’, D’) images showing, colpocephaly i.e., dilated posterior horns (blue arrows), with relatively parallel orientation of the bilateral lateral ventricles suggestive of complete agenesis of the corpus callosum. Second row: Sagittal T2 weighted (E’, F’) and T1 weighted (G’, H’) images showing pachygyria (white arrows) and absent cingulate gyrus (blue arrows). The brainstem and cerebellum are normal (white asterisks). Third Row: Coronal Fluid attenuated inversion recovery (FLAIR) sequences (I’, J’), showing enlarged occipital as well as temporal horns with no periventricular white matter changes. Axial SWI (susceptibility weighted images), (K’, L’) does not show any calcifications. **(C)** EMSA with a labeled DNA probe containing a DMRTA2 consensus binding motif (DmrtA2BS) incubated with (top panel) DMRTA2 WT or DMRTA2^R116P^ proteins produced by TnT and (bottom panel) nuclear extracts from HEK293T cells either non-transfected (/) or transfected with DMRTA2 WT (WT) or DMRTA2 mutated (R116P). Proteins present in the binding reactions are indicated above. Arrows indicate the position of the DNA-protein complexes. “NS” indicates non-specific complexes.

Whether this mutation affects the activity of the protein is an important question to address. Therefore, we first asked whether the mutation affects the subcellular localization of the protein. When overexpressed as a flag fusion in HEK28T3 cells, we found that the mutated protein remains mainly nuclear (data not shown). Since Dmrta2 functions as a DNA-binding transcription repressor, we then tested the ability of the mutated version to bind DNA by electrophoretic mobility shift assays (EMSA). To do so, we incubated the radiolabeled Dmrta2 binding site (Dmrta2BS) probe (Murphy et al., 2007), known to bind the Dmrta2, with either nuclear extracts from HEK293T cells transfected with the expression vector of either the DMRTA2^R116P^ or the WT proteins, or with either the DMRTA2^R116P^ or the WT proteins produced *in vitro* in rabbit reticulocyte lysates. We found that the DMRTA2^R116P^ mutated protein is unable to bind DNA (**Fig. 6C**). With this *in vitro* evidence of the plausible pathogenicity of the *DMRTA2R116PVUS*, this variant can thus be re-classified as likely-pathogenic following the updated ACMG criteria (PS3, PM2, PP1, PP3, and PP4). Together, these results suggest that DMRTA2^R116P^ causes cortical malformations by affecting its DNA binding properties.

### Dmrta2 interacts with the NurD complex and Zfp423

How Dmrta2 represses gene expression is unknown. To identify Dmrta2 interacting partners, we generated transgenic mice expressing Dmrta2 with an N-terminal 2 X HA epitope using the CRISPR/Cas9 gene-editing system (*Dmrta2^2XHA^*). **Fig. 7A** shows RT-qPCR, western blot, and immunostaining experiments demonstrating that the tagged *Dmrta2* allele is expressed in the telencephalon of *Dmrta2^2XHA^* embryos. We then performed rapid immunoprecipitation followed by RIME experiments using E12.5 dissected dorsal cortices of *Dmrta2^2XHA^* and WT control embryos. As this method uses formaldehyde fixation to stabilize proteins complexes, it is particularly suited to study chromatin and transcription factor complexes (Mohammed et al., 2016). Among the 263 proteins that selectively copurified with Dmrta2-HA (**Fig. 7B**), the major protein present in the *Dmrta2^2XHA^* sample was Dmrta2 in two independent replicates. Other members of the DmrtA family, such as Dmrt3 and Dmrta1, were also efficiently recovered, which is expected given the ability of DmrtA proteins to bind DNA, forming homo-and heterodimers, trimers, or tetramers (Murphy et al., 2007, 2015). Interestingly, several components of the nucleosome remodeling and deacetylase NuRD complex (including CHD4, HDAC2, MBD3 among others), and the vertebrate-specific zinc finger transcription factor Zfp423 (also termed OLF/EBF associated-zinc finger protein OAZ) and its homolog Zfp521 that interact with the NuRD complex (Alcaraz et al., 2006; Harder et al., 2014; Nitarska et al., 2016; Li et al., 2021) and regulate neurogenesis (Cheng et al., 2007; Kamiya et al, 2011; Shen et al., 2011; Ohkubo et al., 2014; Shahbazi et al., 2016; Massimino et al., 2018) were also selectively enriched in the *Dmrta2^2XHA^* samples (**Fig. 7C-E**). Among the other proteins recovered selectively in the *Dmrta2^2XHA^* sample was the zinc-finger protein Zfp462 which regulates neural lineage specification by targeting the H3K9-specific histone methyltransferase complex G9A/GLP to silence meso-endodermal genes (Yelagandula et al., 2023) and the E3 ubiquitin ligases BRE1a and BRE1b that regulate the cell cycle and differentiation of neural precursor cells (NPCs) through monoubiquitynation of histone H2B (Ishino et al., 2014).

**Figure 7.**
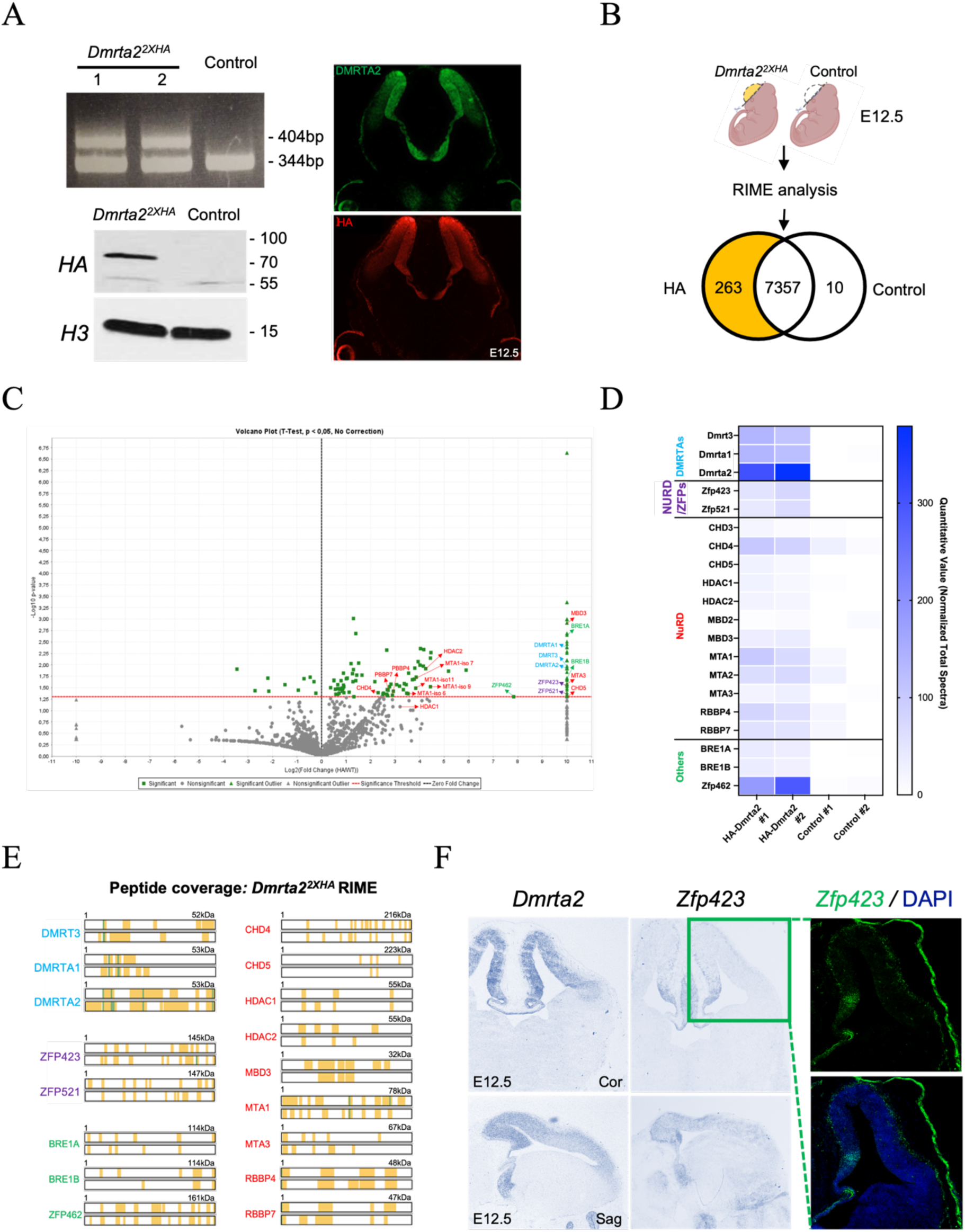
RIME-purified Dmrta2 complexes isolated from dissected cortical tissue contain multiple subunits of NuRD and the NuRD interacting Zfp423 protein. **(A)** Left: RT-qPCR using cDNA prepared from dissected cortices of E12.5 embryos with primers flanking the inserted 2XHA sequence, showing that the tagged allele is detectable in the two F0 *Dmrta2^2XHA^* mice (clones # 1 and # 2) and is absent in WT mice. A Western blot is shown below demonstrating that the HA-tagged protein is expressed in *Dmrta2^2xHA^*mice. Right: immunostainings showing that the HA-tagged allele is expressed similarly to Dmrta2 in the telencephalon. **(B)** Schematic diagram of the experiment performed. Cortices were dissected from E12.5 *Dmrta2^2xHA^*and control embryos and subjected to RIME analysis. Venn diagrams with the number of the differentially co-purified proteins identified in the cortex of *Dmrta2^2xHA^* and control embryos are shown. **(C)** A Volcano plot showing the identified Dmrta2 interacting proteins identified using RIME. All statistically validated proteins are represented by green points, while grey points represent background binding proteins. An arbitrary -log10 p-value of 10 was attributed to the outliers, corresponding to proteins detected only in the HA-Dmrta2 experimental conditions. Proteins of interest are indicated with name (n=3 for each condition). Dmrta2 and other DMRTA family members are highlighted in bright blue, NuRD members in red, and NuRD complex interacting zinc-finger proteins in orange. **(D)** A heatmap showing the significantly enriched proteins identified by RIME and their normalized total spectral count across indicated samples (n=2 for each condition). **(E)** Peptide coverage for some of the Dmrta2 interacting proteins was identified with a significance level of P < 0.05 in *Dmrta2^2XHA^* embryos (n=2 for each condition). Yellow bars represent regions of the full-length protein sequence where peptides were identified. **(F)** ISH on coronal and sagittal sections of the brain of an E12 WT embryo for *Dmrta2* and *Zfp423*, and immunofluorescence of coronal sections with Zfp423 antibodies.

Zfp521 can act as a repressor or activator depending on the cellular context (Scicchitano et al., 2019; Chiarella et al., 2021). During the differentiation of embryonic stem cells into neural progenitors, Zfp521 has been shown to act mainly as an activator of neural genes (Kamiya et al., 2011). Zfp462 has been shown to repress mesoendodermal genes during neural differentiation of embryonic stem cells but does not affect *Pax6* upregulation (Yelagandula et al., 2023). Therefore, and based on our data showing that Dmrta2 complexes contain multiple NuRD subunits, we focused on the NurD-interacting Zfp423 protein as a possible Dmrta2 interacting partner potentially involved in its repressive function. By *in situ* hybridization and immunostaining, we found as previously reported (Cheng et al., 2007; Massimino et al., 2018) that *Zfp423* is expressed in cortical progenitors, the highest expression being detected in the hem region (**Fig. 7F**). To further investigate the ability of Dmrta2 to form a complex with Zfp423, constructs encoding *Zfp423* and Flag-tagged *Dmrta2* were cotransfected in HEK293T cells. In co-immunoprecipitation (Co-IP) assays, results obtained indicated that Flag-Dmrta2 interacts with Zfp423 (**Fig. 8A**) and revealed that the DM domain of Dmrta2 is required for the interaction (**Fig. 8B**). As Zfp423 interacts with the NuRD complex, its interaction with Dmrta2 may contribute to its repressive properties. To test this hypothesis, *Zfp423* was co-transfected at increasing doses with a fixed dose of Flag*-Dmrta2* in HEK293T cells. Co-IP assays were then performed using an anti-Flag antibody followed by western blotting using antibodies for Flag-Dmrta2, Zfp423, and NuRD complex components. Despite some background binding observed with protein G magnetic beads alone used as a negative control, the results obtained suggest that Dmrta2 can immunoprecipitate some NuRD subunits (i.e HDAC1 and MBD3, compare levels in lines 1 and 5). With increasing doses of Zfp423, as expected, increasing binding of Zfp423 was observed. Interestingly, increased binding of Zfp423 to Dmrta2 appears to increase the amount of coimmunoprecipitated HDAC1/2 and MBD3 (**Fig. 8C and Extended data Fig. 8-1**). These results suggest that Dmrta2 can interact with NuRD subunits and that Zfp423 interaction enhances its association with NuRD components.

**Figure 8.**
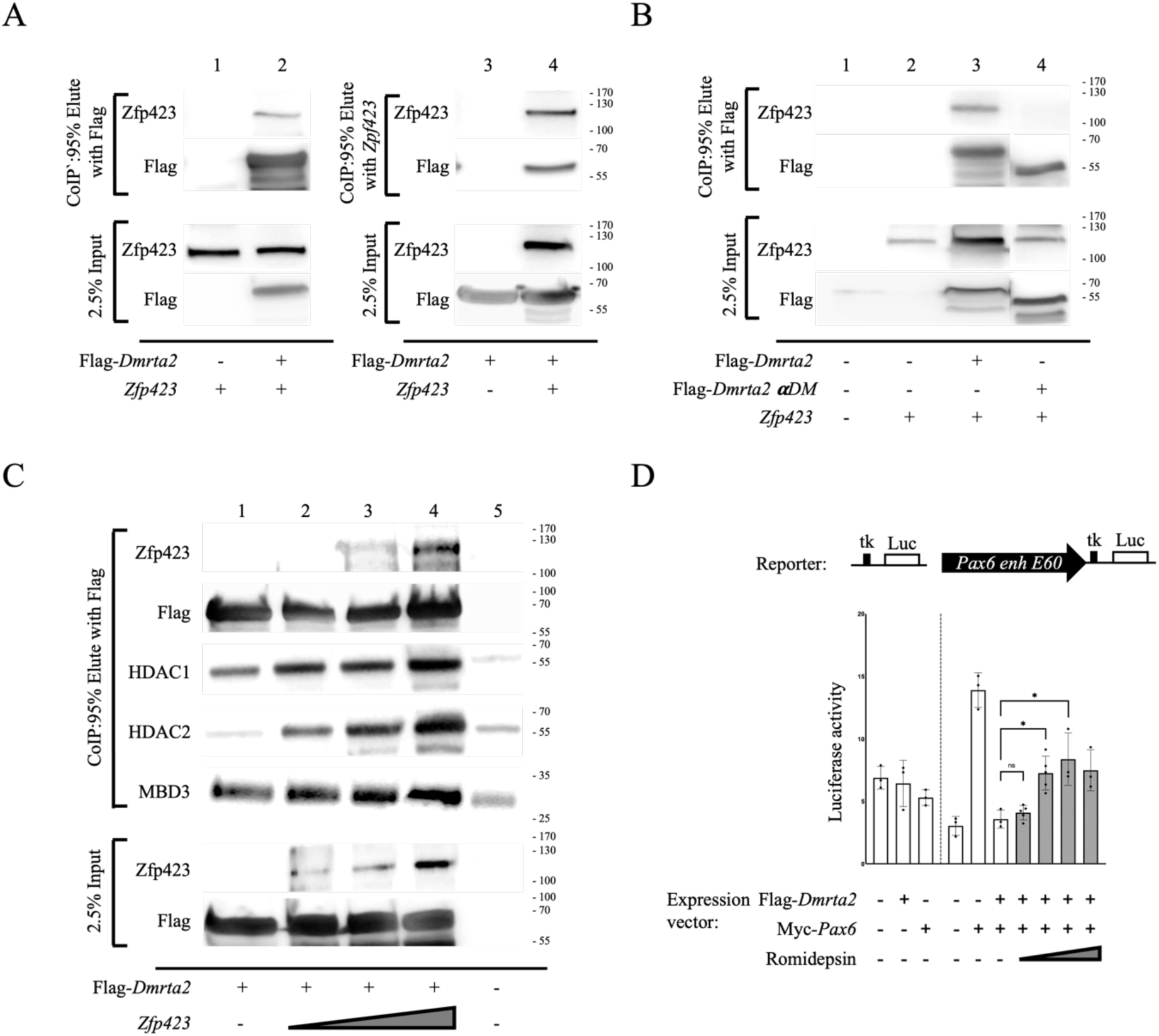
The Dmrta2-Pax6 interaction requires the recruitment of the HDAC-NuRD complex by Zfp423. **(A, B)** Co-immunoprecipitation assays as indicated with *Zfp423,* Flag*-Dmrta2,* and a Flag*-Dmrta2* mutant lacking the DM domain (Flag-*Dmrta2 **a**DM*) overexpressed in HEK293T cells as indicated. Note in A that Flag-Dmrta2 pulled down Zpf423 (line 1-2) and that, conversely, Zpf423 co-precipitated Flag-Dmrta2 (line 3-4). Note in B that Flag-Dmrta2 ***α***DM does not pull down Zfp423 (line 4). **(C)** Co-immunoprecipitation assays as indicated with HEK293T cells transfected with a Flag*-Dmrta2* expression construct, alone or together with increasing doses of a *Zfp423* expression construct. Note that Dmrta2 immunoprecipitates some NuRD subunits and that the binding of Zfp423 to Dmrta2 appears to increase the amount of coimmunoprecipitated HDAC1/2 and MBD3. **(D)** Reporter assays in P19 cells transfected with a *Pax6 E60* tk-luciferase reporter vector, or an ‘empty’ tk-luciferase reporter vector as indicated, together with a Myc*-Pax6* expression vector and/or a Flag*-Dmrta2* expression vector as indicated, in the absence (white bars) or presence of increasing doses of the HDAC1 inhibitor romidepsin (grey bars). Not that romidepsin partially blocks the ability of Dmrta2 to repress the activity of the *Pax6 E60* enhancer. *P < 0.05.

Finally, to address a potential role for Zfp423 in Dmrta2 repressive function, we performed reporter assays using the *Pax6 E60*-luc reporter construct in P19 cells transfected as above with a *Pax6* expression vector to stimulate *E60* enhancer activity, in the presence or absence of co-transfected Flag*-Dmrta2*. 24h after transfection, cells were treated with the class I HDAC inhibitor romidepsin, which is known for its selective inhibition of HDAC 1, 2, 3, and 8 (Petrich and Nabhan, 2016; Mayr et al., 2021). Results obtained showed that romidepsin partially blocks the ability of Dmrta2 to repress the activity of the *Pax6 E60* enhancer (**Fig. 8D**). Together, these data provide the first evidence that Dmrta2 functions as a transcriptional repressor through the recruitment of the NuRD complex. They also suggest that Zfp423 collaborate with Dmrta2 to enhance its repressive properties.

## DISCUSSION

In this study, we provided an extensive analysis of how Dmrta2 and Pax6, two transcription factors expressed in the cortical ventricular zone in opposite gradients, interact to control the spatiotemporal identity of neural progenitors and regulate regional neurogenesis. We show that *Pax6* is already upregulated by the loss of *Dmrta2* at E9.5 before the disruption of Wnt signaling, in accordance with its identification as a Dmrta2 direct target (Konno et al., 2019). *Pax6* promotes neurogenesis while *Dmrta2* maintains neural progenitors in the cell cycle (Scardigli et al., 2003; Sansom et al., 2009; Young et al., 2017). We found that the absence of one allele of *Pax6* partially rescues the reduction of the size of the cortex observed in *Dmrta2^-/-^* embryos due to premature neuronal differentiation. This is in accordance with the observation that the siRNA knockdown of *Pax6* expression in medial cortical cells electroporated with siRNAs targeting *Dmrt3* and *Dmrta2* rescues their neurogenic phenotype. This observation further suggests that Dmrt factors regulate region-dependent neurogenic properties of neural progenitors by establishing a lateral ^high^/medial ^low^ gradient of *Pax6* in the developing dorsal telencephalon, as proposed by Konno et al.(2019).

Both Pax6 (Toresson et al., 2000, Yun et al., 2001; Carney et al., 2009) and Dmrta2 (Desmaris et al., 2018) have been reported to be required for defining the dorsal telencephalic compartment by repressing Gsx2. Here we show that the subpallial markers Gsx2 is ectopically expressed in the abortive cortical primordium of *Pax6; Dmrta2* double KO embryos, much more robustly than in single KO embryos, indicating that they cooperate in maintaining cortical identity in dorsal telencephalic progenitors. The combined loss of *Pax6* and *Emx2* (Muzio et al., 2002) and of *Dmrta2* and *Emx2* (Desmaris et al., 2018) also results in the ectopic expression of ventral-specific markers in the dorsal telencephalon. Early telencephalon dorsoventral patterning and the maintenance of the identity of cerebral cortical progenitors appear thus to be under the tight control of multiple cortical factors that cooperate to repress the expression of ventral telencephalic determinants such as Gsx2.

*Pax6* is expressed in a rostrolateral ^high^/caudomedial ^low^ gradient, opposite to that of *Dmrta2*. *Pax6* regulates dorsoventral patterning in the telencephalon. Within the lateral telencephalon, *Pax6* contributes to the restriction of medial cortical fate (Muzio et al., 2002; Godbole et al., 2017). *Pax6* also promotes a rostral fate in cortical progenitors. In small eye mutants that lack functional Pax6 protein, caudal visual areas are expanded (Bishop et al., 2000; 2002). The cortex-specific deletion or overexpression of *Pax6* has been also shown to reduce S1 area size (Manuel et al., 2007; Zembrzycki et al., 2013). The study by Konno et al. (2019), suggested that the graded expression of Dmrt factors provides positional information and thus confers identity to cortical progenitors by suppressing *Pax6* transcription and establishing its lateral ^high^/medial ^low^ expression gradient. Our findings that 1) medial cortical hem fate can be rescued in *Dmrta2* hetero- and homozygous mutant embryos by the reduction of *Pax6* and 2) that *Pax6* overexpression rescues the expansion of V1 area observed in *Dmrta2* overexpressing transgenics provide first experimental support to this model.

How mechanistically Dmrta2 acts to control gene expression in cortical progenitors remains unknown. To answer this question, we first performed reporter gene assays in P19 cells using the *Pax6 E60* enhancer. Given the low activity of this enhancer in P19 cells, those assays were performed in the presence of *Pax6*, as previous studies reported that Pax6 autoregulates its expression (Aota et al., 2003; Manuel et al., 2007, 2015) and that Pax6 binds to this enhancer region (Sun et al., 2015). In accordance with these studies, our results show that Pax6 increases luciferase activity using this *E60* enhancer-driven reporter construct. However, this is not in agreement with the observation that this *E60* enhancer remains active in *Pax6*^sey/sey^ mutants (McBride et al., 2011). The reason for this discrepancy remains unclear. Our results indicate that Dmrta2 counteracts the activation by Pax6 of this *E60* enhancer-driven reporter and that the zinc finger motifs of Dmrta2 that are required for DNA binding (Murphy et al, 2015) are essential for this repression. These results suggest that Dmrta2 acts as a DNA-binding repressor on the *Pax6* locus, as it appears to be the case on the repressor of neurogenesis *Hes1* and on the *Gsx2* genomic loci (Young et al., 2017; Desmaris et al., 2018).

This is also further supported by the identification of the human DMRTA2^R116P^point mutation reported here that, to our knowledge, is the only mutation in DMRTA2 suggested to play a role in microcephaly by affecting its ability to bind DNA. The DMRTA2^R116P^ mutation corresponds to DMRT1^R/K122^ involved in DNA phosphate backbone contacts. The fact that the change is to a proline, a helix-breaking residue, and occurs close to DMRT1^R12*3*^ that makes base-specific contacts likely explains the observed disruption of DNA binding. An animal model of the DMRTA2^R116^ mutation may help to elucidate the cause of the phenotype observed in this family.

Besides the identification of its direct targets, another critical step in the understanding of the mode of action of a transcription factor is the identification of the protein complexes in which it acts. Our data suggest that Dmrta2 interacts with the NuRD complex and with the multi-zinc finger Zfp423 and related Zfp521 proteins that can also interact with NurD components (Harder et al., 2014) and that the association of Dmrta2 with Zfp423 enhances its repressive properties. Both proteins are expressed in cortical progenitors (Kamiya et al., 2011; Massimino et al., 2018). They act as scaffolds or as transcription factors, cooperating with multiple regulatory molecules. They both play important roles in neural development. The importance of Zfp521 for neural development is demonstrated by the fact that it is essential and sufficient for driving the intrinsic neural differentiation of mouse ES cells (Kamiya et al., 2011; Shahbazi et al., 2016). ZNF423 mutations are associated with Joubert Syndrome, a ciliopathy causing cerebellar vermis hypoplasia and ataxia. Null Zfp423 mutants develop cerebellar malformations and hindbrain choroid plexus hypoplasia (Cheng et al., 2007; Casoni et al., 2017, 2020). In the forebrain, the corpus callosum is absent and the hippocampus is reduced. The cortex is thinner than the controls due to reduced proliferation. When Zfp423 is overexpressed in the cortex of E13.5 mouse embryos, it increases the number of electroporated cells positive for the neuronal marker *Tubb3* at the expense of mitotically active PAX6^+^ radial glia cells (Alcaraz et al., 2006; Massimino et al., 2018). Whether these changes in cell proliferation and differentiation upon manipulation of Zpf423 expression are linked to its interaction with Dmrta2 or, alternatively, to its ability to modulate other signaling pathways, like the SMAD/BMP, NOTCH, and SHH pathways (Hata et al., 2000; Ku et al., 2006; Masserdotti et al., 2010; Hong and Hamilton, 2016) remains to be elucidated.

## EXTENDED DATA

**Extended data Fig. 1-1.**
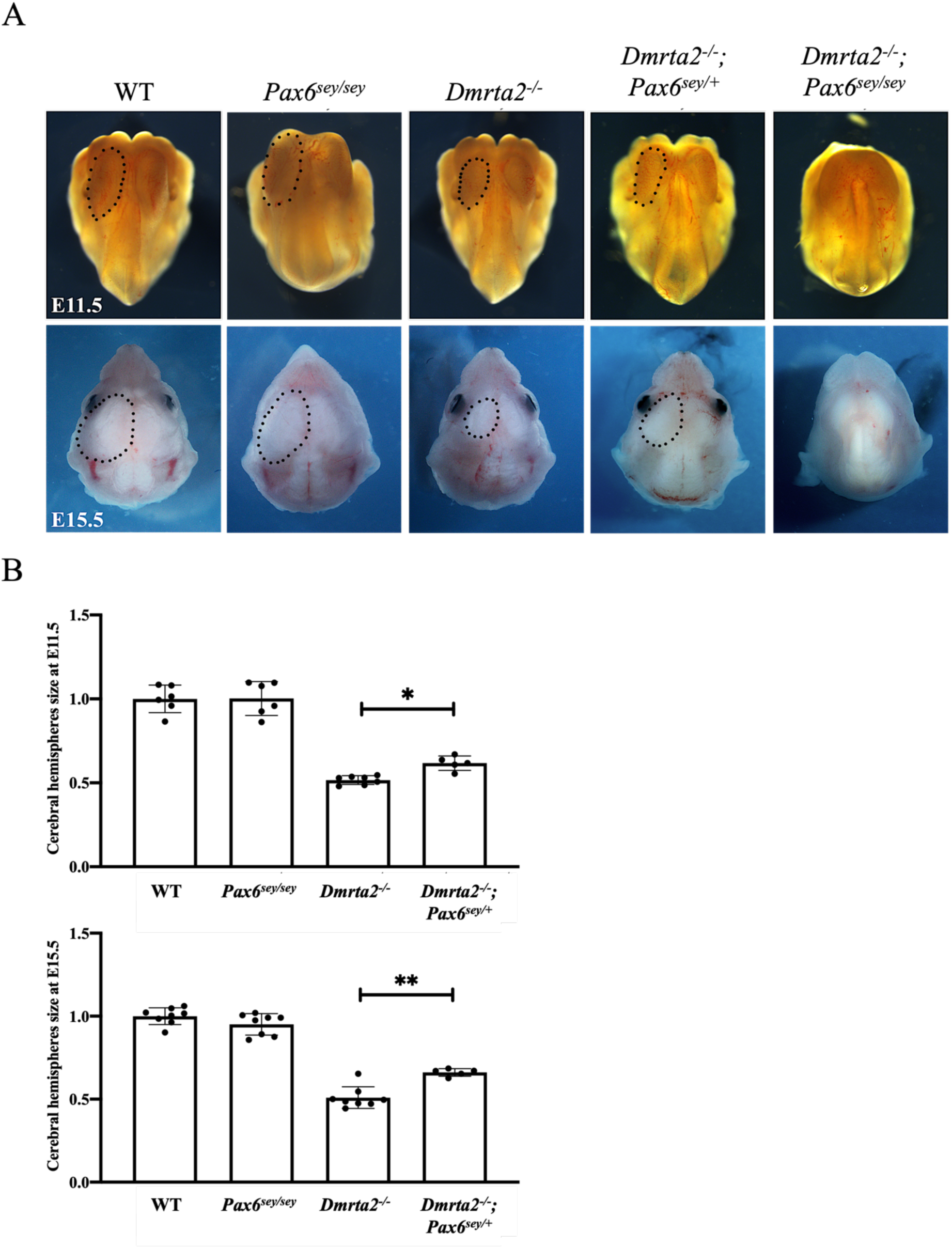
The rescue of the growth of telencephalic vesicles upon the loss of one *Pax6* allele in *Dmrta2*^-/-^ embryos is detectable in early embryos. **(A)** Dorsal views of the brain of E11.5 and E15.5 embryos of the indicated genotype. (B) Graphs representing the surface area of E11.5 and E15.5 cerebral hemispheres compared to WT set to 1.*P < 0.05, **P < 0.01.

**Extended data Fig. 5-1.**
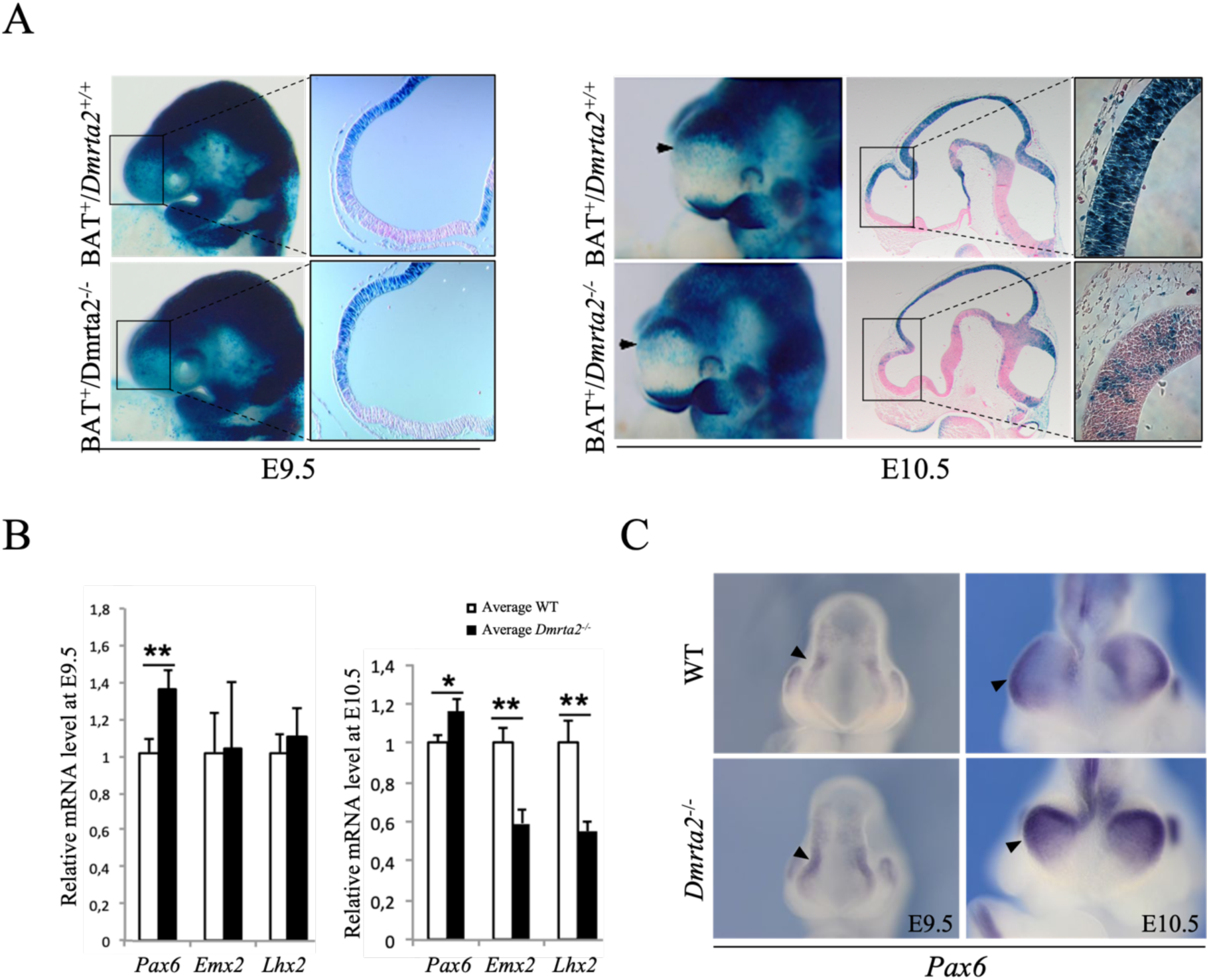
*Pax6* is already upregulated by the loss of *Dmrta2* at E9.5 before the Wnt signaling pathway appears to be affected. **(A)** Whole-mount X-gal staining on BAT^+^/*Dmrta2^+/+^* and BAT^+^/*Dmrta2^-/-^* embryos at E9.5 and E10.5. Note on the whole-mount views (arrows) and on the sections and magnification views shown on the left that the staining detected in BAT^+^/*Dmrta2^-/-^* embryos is similar to that of BAT^+^/*Dmrta2^+/+^* embryos at E9.5, but is reduced at E10.5. **(B)** RT–qPCR analysis of *Pax6, Emx2,* and *Lhx2* expression in the cortex of *Dmrta2^-/-^* and WT embryos. Results are normalized to the level of expression in the cortex of WT embryos. Note that *Pax6* is already upregulated in the cortex of *Dmrta2^-/-^* at E9.5, which is not the case of *Emx2* and *Lhx2* whose deregulation is only observed from E10,5. Error bars indicate SDs of at least 3 independent experiments. *P <0.05, **P < 0.01. **(C)** Whole mount ISH analysis of *Pax6* expression shows that its upregulation (arrow) in *Dmrta2^-/-^* embryos can be already seen from E9.5.

**Extended data Fig. 5-2.**
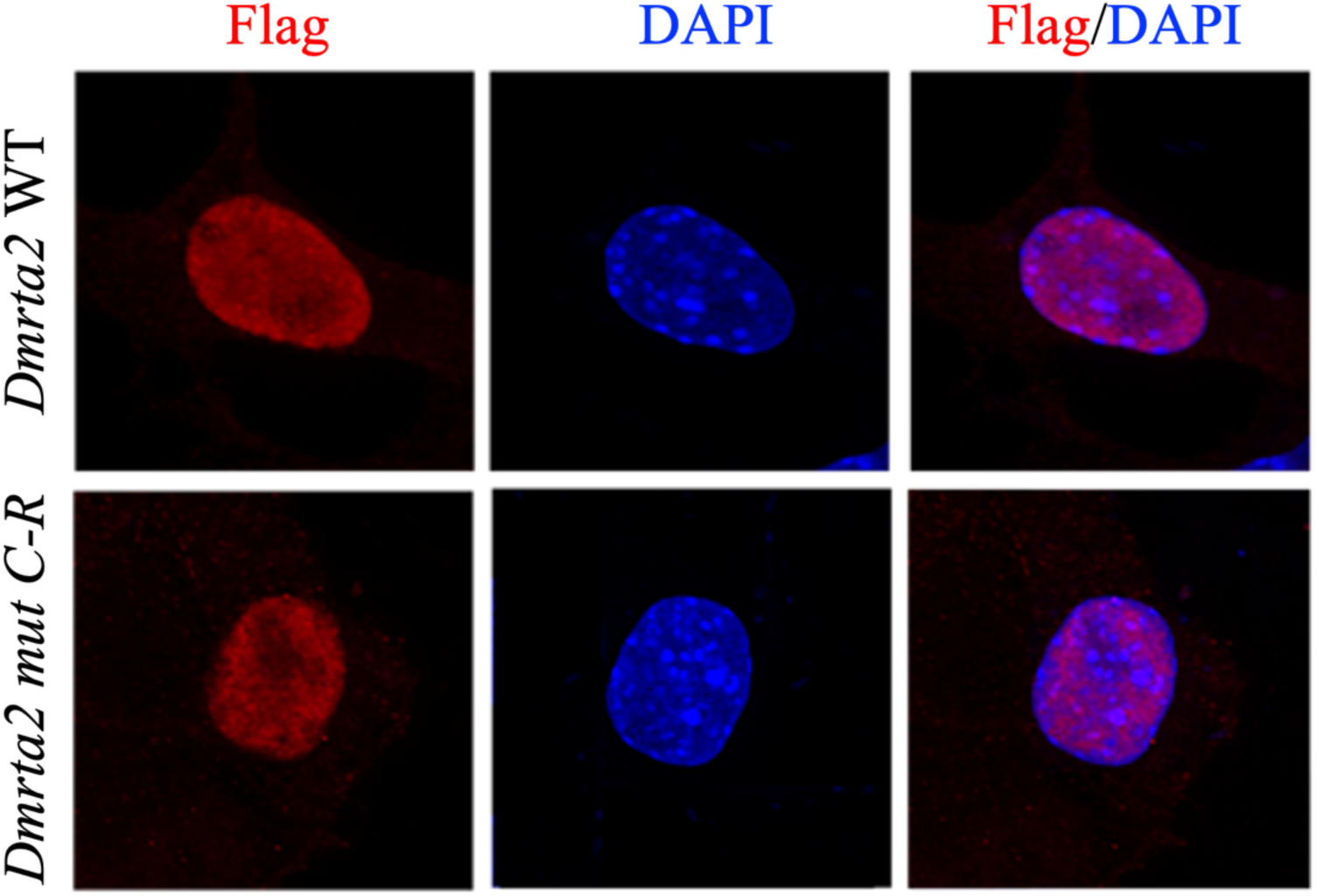
The Flag-*Dmrta2 mut C-R is* detected in the nucleus, as the WT protein in transfected P19 cells. Flag immunostaining and DAPI staining are shown, together with a merged image.

**Extended data Table 6-1:**
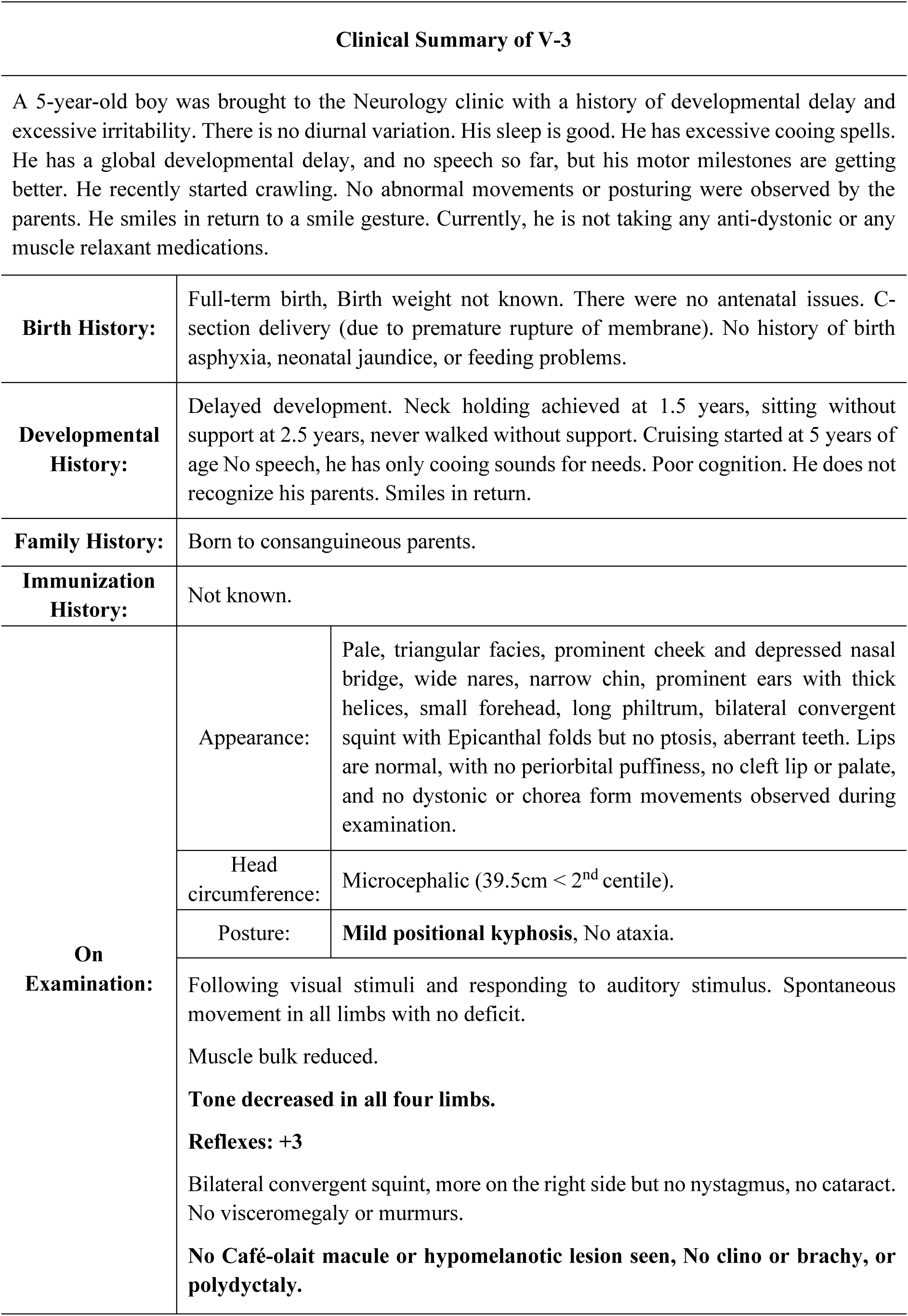
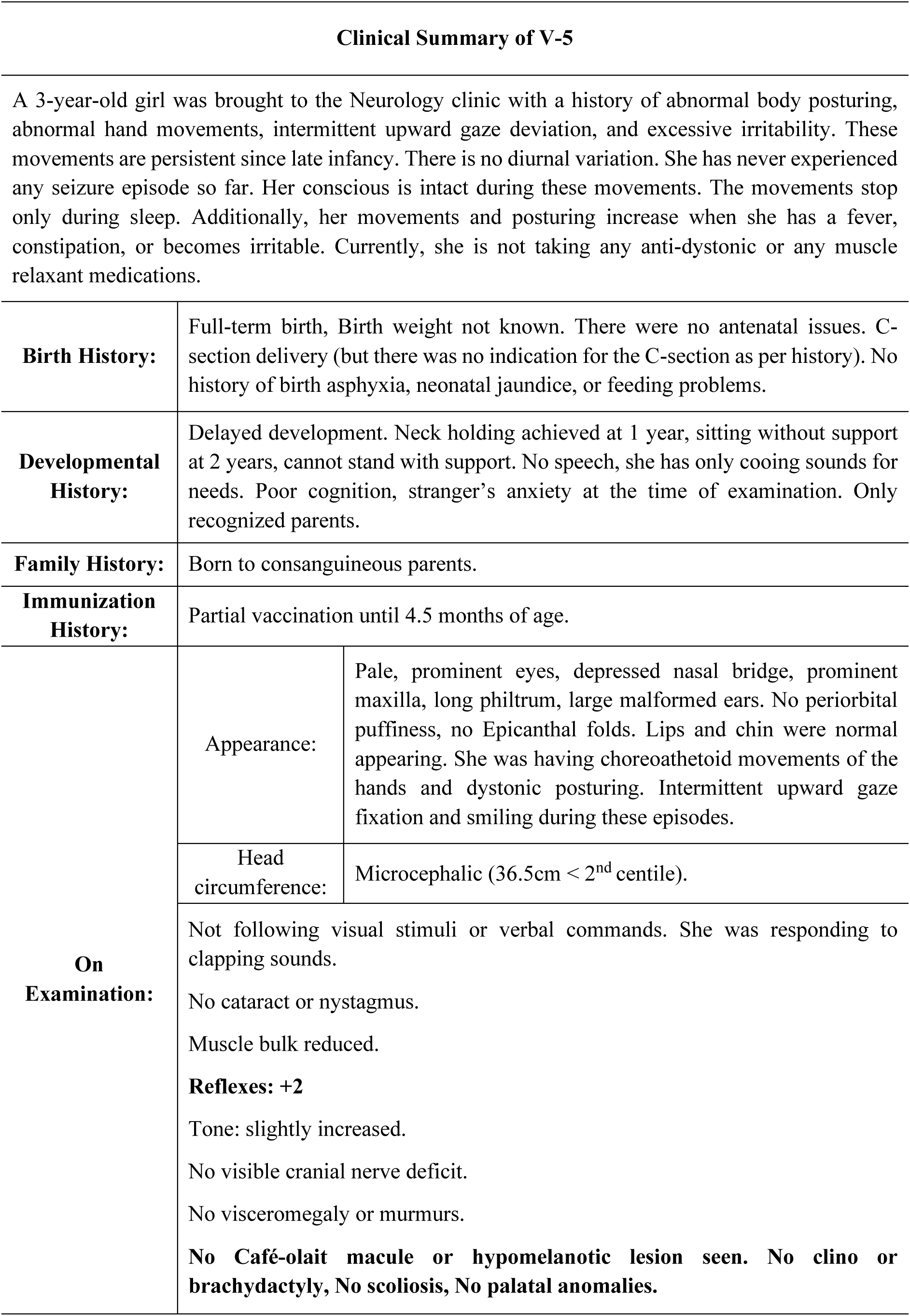
Clinical evaluation of V-3 and V-5 patients.

**Extended data Fig. 8-1.**
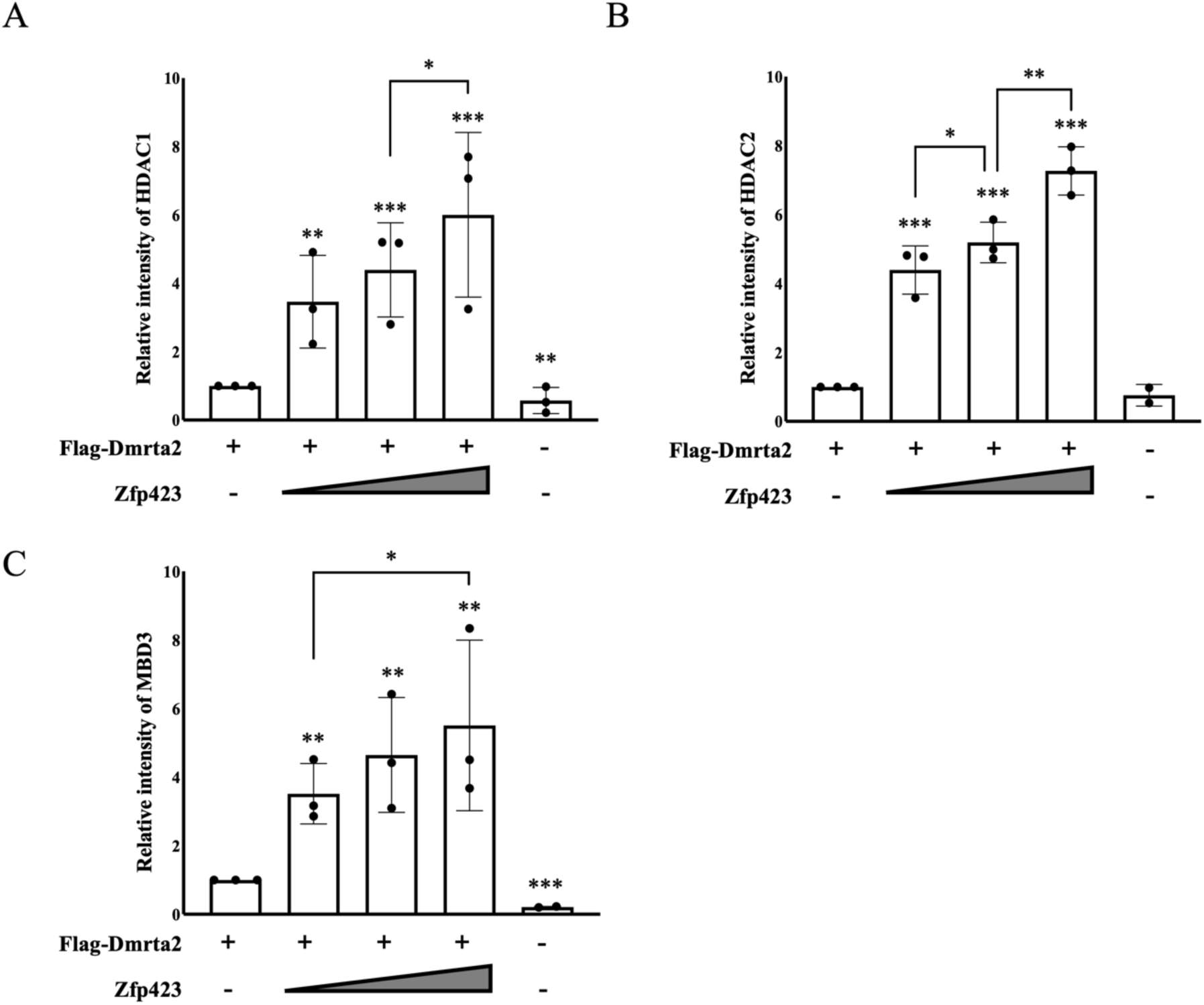
Densitometric quantification of Western blot results of HDAC1 (A), HDAC2 (B), and MBD3 (C) from Figure 8C. For each comparison, the value for the expression levels in cells transfected with Flag-*Dmrta2* alone is set as 1. The data are presented as a comparison between these baseline cells and those transfected with additional constructs.* P < 0.05, ** P < 0.01, *** P < 0.001.

## Notes

### Competing Interest Statement

The authors have declared no competing interest.

## REFERENCE

Alcaraz WA, Gold DA, Raponi E, Gent P, Conception D, Hamilton BA (2006) Zfp423 controls proliferation and differentiation of neural precursors in cerebellar vermis formation. Proc Natl Acad Sci U S A 103:19424–19429.

Aota SI, Nakajima N, Sakamoto R, Watanabe S, Ibaraki N, Okazaki K (2003) Pax6 autoregulation mediated by direct interaction of Pax6 protein with the head surface ectoderm-specific enhancer of the mouse Pax6 gene. Dev Biol: 257, 1–13.

Berger J, Berger S, Tuoc TC, D’Amelio M, Cecconi F, Gorski JA, Jones KR, Gruss P, Stoykova A (2007) Conditional activation of Pax6 in the developing cortex of transgenic mice causes progenitor apoptosis. Development 134:1311–1322.

Bishop KM, Goudreau G, O’Leary DD (2000) Regulation of area identity in the mammalian neocortex by Emx2 and Pax6. Science 288: 344–349.

Bishop KM, Rubenstein JL, and O’Leary DD (2002) Distinct actions of Emx1, Emx2, and Pax6 in regulating the specification of areas in the developing neocortex. J Neurosci 22: 7627–7638.

Cadwell CR, Bhaduri A, Mostajo-Radji MA, Keefe MG, Nowakowski TJ (2019) Development and arealization of the cerebral cortex. Neuron 103: 980–1004.

Carney RS, Cocas LA, Hirata T, Mansfield K, and Corbin JG (2009) Differential regulation of telencephalic pallial–subpallial boundary patterning by Pax6 and Gsh2. Cereb Cor 19: 745–759.

Casoni F, Croci L, Bosone C, Hawkes R, Warning S, Consalez GG (2017) Zfp423/ZNF423 regulates cell cycle progression, the mode of cell division and the DNA-damage response in Purkinje neuron progenitors. Development 144: 3686–3697.

Casoni F, Croci L, Vincenti F, Podini P, Riba M, Massimino L, Cremona O, Consalez GG (2020) ZFP423 regulates early patterning and multiciliogenesis in the hindbrain choroid plexus. Development 147: dev190173.

Cheng LE, Zhang J, and Reed RR (2007) The transcription factor Zfp423/OAZ is required for cerebellar development and CNS midline patterning. Devl Biol 307:43–52.

Chiarella E, Aloisio A, Scicchitano S, Todoerti K, Cosentino EG, Lico D, Neri A, Amodio N, Mandy Bond H, Mesuraca M (2021) ZNF521 Enhances MLL-AF9-Dependent Hematopoietic Stem Cell Transformation in Acute Myeloid Leukemias by Altering the Gene Expression Landscape. Int J Mol Sci 22: 10814.

De Clercq S, Keruzore M, Desmaris E, Pollart C, Assimacopoulos S, Preillon J, Ascenzo S, Matson CK, Lee M, Nan X, Li M, Nakagawa Y, Hochepied T, Zarkower D, Grove EA, Bellefroid EJ (2018) DMRT5 together with DMRT3 directly controls hippocampus development and neocortical area map formation. Cereb Cortex 28: 493–509.

Desmaris E, Keruzore M, Saulnier A, Ratié L, Assimacopoulos S, De Clercq S, Henningfeld K, Zarkower D, Mallamaci A, Campbell K, Pieler T, Li M, Grove EA, Bellefroid EJ (2018) DMRT5, DMRT3, and EMX2 cooperatively repress GSX2 at the pallium–subpallium boundary to maintain cortical identity in dorsal telencephalic progenitors. J Neurosci 38: 9105–9121.

Dignam JD, Lebovitz RM, Roeder RG (1983) Accurate transcription initiation by RNA polymerase II in a soluble extract from isolated mammalian nuclei. Nucleic Acids Res 11:1475– 89.

Godbole G, Roy A, Shetty AS, Tole S (2017) Novel functions of LHX2 and PAX6 in the developing telencephalon revealed upon combined loss of both genes. Neural Development 12: 1–8.

Hamasaki T, Leingärtner A, Ringstedt T, O’Leary DD (2004) EMX2 regulates sizes and positioning of the primary sensory and motor areas in neocortex by direct specification of cortical progenitors. Neuron 43: 359–372.

Harder L, Puller AC, Horstmann MA (2014) ZNF423: transcriptional modulation in development and cancer. Mol Cell Oncol 1: e969655.

Hata A, Seoane J, Lagna G, Montalvo E, Hemmati-Brivanlou A, and Massagué J (2000) OAZ uses distinct DNA-and protein-binding zinc fingers in separate BMP-Smad and Olf signaling pathways. Cell 100: 229–240.

Hill RE, Favor J, Hogan BL, Ton CC, Saunders GF, Hanson IM, Prosser J, Jordan T, Hastie ND, van Heyningen V (1991) Mouse small eye results from mutations in a paired-like homeobox-containing gene. Nature, 354:522–525.

Hong CJ, Hamilton BA (2016) Zfp423 regulates sonic hedgehog signaling via primary cilium function. PLoS Genet 12: e1006357.

Ishino Y, Hayashi Y, Naruse M, Tomita K, Sanbo M, Fuchigami T, Fujiki R, Hirose k, Toyooka Y, Fujimori t, Ikenaka K, Hitoshi S (2014) Bre1a, a histone H2B ubiquitin ligase, regulates the cell cycle and differentiation of neural precursor cells. J Neurosci 34: 3067–3078.

Kamiya D, Banno S, Sasai N, Ohgushi M, Inomata H, Watanabe K, Kawada M, Yakura R, Kiyonari H, Nakao K, Martin L, Martin Jakt L, Nishikaa S-I, Sasai Y (2011) Intrinsic transition of embryonic stem-cell differentiation into neural progenitors. Nature 470: 503–509.

Kircher M, Witten DM, Jain P, O’roak BJ, Cooper GM, Shendure J (2014) A general framework for estimating the relative pathogenicity of human genetic variants. Nat Genet 46: 310–315.

Konno D, Iwashita M, Satoh Y, Momiyama A, Abe T, Kiyonari H, Matsuzaki F. (2012). The mammalian DM domain transcription factor Dmrta2 is required for early embryonic development of the cerebral cortex: PlosOne 7:e46577.

Konno D, Kishida C, Maehara K, Ohkawa Y, Kiyonari H, Okada S, Matsuzaki F (2019) Dmrt factors determine the positional information of cerebral cortical progenitors via differential suppression of homeobox genes. Development 146: dev174243.

Ku M, Howard S, Ni W, Lagna G, Hata A (2006) OAZ regulates bone morphogenetic protein signaling through Smad6 activation. J Biol Chem 281: 5277–5287.

Lee SM, Tole S, Grove EA, McMahon AP (2000) A local Wnt-3a signal is required for development of the mammalian hippocampus. Development 127:457–467.

Li Z, Fu X, Wu W, Liu Z, Chen Z, Zhou C, Liu Y, Kuang M, Sun F, Xiao F, Huang Y, Zhang X, Fan S, Huang X, Zheng G, Chen J, Hou Y (2021) Zfp521 is essential for the quiescence and maintenance of adult hematopoietic stem cells under stress. Iscience 24:102039.

Manuel M, Georgala PA, Carr CB, Chanas S, Kleinjan DA, Martynoga B, Mason JO, Molinek M, Pinson J, Pratt T, Quinn JC, Simpson TI, Tyas DA, van Heyningen V, Price DJ (2007) Controlled overexpression of Pax6 *in vivo* negatively autoregulates the Pax6 locus, causing cell-autonomous defects of late cortical progenitor proliferation with little effect on cortical arealization. Development 134: 545–555.

Manuel MN, Mi D, Mason JO, Price DJ (2015) Regulation of cerebral cortical neurogenesis by the Pax6 transcription factor. Front Cell Neurosci 9: 70.

Maretto S, Cordenonsi M, Dupont S, Braghetta P, Broccoli V, Hassan AB, Volpin D, Bressan GM, Piccolo S (2003) Mapping Wnt/β-catenin signaling during mouse development and in colorectal tumors. Proc Natl Aca Sci USA100: 3299–3304.

Masserdotti G, Badaloni A, Song YS, Croci L, Barili V, Bergamini G, Vetter M, Consalez GG (2010) ZFP423 coordinates Notch and Bone Morphogenetic Protein (BMP) signaling, selectively upregulating Hes5 gene expression. J Biol Chem 285: 30814–30824.

Massimino L, Flores-Garcia L, Di Stefano B, Colasante G, Icoresi-Mazzeo C, Zaghi M, Hamilton BA, Sessa A (2018) TBR2 antagonizes retinoic acid dependent neuronal differentiation by repressing Zfp423 during corticogenesis. Dev Biol 434: 231–248.

Mayr C, Kiesslich T, Erber S, Bekric D, Dobias H, Beyreis M, Ritter M, Jäger T, Neumayer B, Winkelmann P, Klieser E, Neureiter D (2021) HDAC Screening Identifies the HDAC Class I Inhibitor Romidepsin as a Promising Epigenetic Drug for Biliary Tract Cancer. Cancers, 13: 3862.

McBride DJ, Buckle A, van Heyningen V, Kleinjan DA (2011) DNaseI hypersensitivity and ultraconservation reveal novel, interdependent long-range enhancers at the complex Pax6 cis-regulatory region. PloS One 6: e28616

Meier F, Brunner AD, Koch S, Koch H, Lubeck M, Krause M, Goedecke N, Decker J, Kosinki T, Park MA, Bache N, Hoerning O, Cox j, Räther O, Mann M (2018) Online parallel accumulation–serial fragmentation (PASEF) with a novel trapped ion mobility mass spectrometer. Mol Cell Proteomics, 17: 2534–2545.

Mohammed H, Taylor C, Brown GD, Papachristou EK, Carroll JS, D’santos CS (2016) Rapid immunoprecipitation mass spectrometry of endogenous proteins (RIME) for analysis of chromatin complexes. Nature Protoc 11: 316–326.

Monuki ES, Porter FD, Walsh CA (2001) Patterning of the dorsal telencephalon and cerebral cortex by a roof plate-Lhx2 pathway. Neuron. 32:591–604.

Murphy MW, Zarkower D, Bardwell VJ. (2007). Vertebrate DM domain proteins bind similar DNA sequences and can heterodimerize on DNA. BMC Mol Biol 8: 1–14.

Murphy MW, Lee JK, Rojo S, Gearhart MD, Kurahashi K, Banerjee S, Loeuille G-A, Bashamboo A, McElreavey K, Zarkower D, Aihara H, Bardwell VJ (2015) An ancient protein-DNA interaction underlying metazoan sex determination. Nat Struct Mol Biol 22: 442–451.

Muzio L, Di Benedetto B, Stoykova A, Boncinelli E, Gruss P, Mallamaci A (2002) Emx2 and Pax6 control regionalization of the pre-neuronogenic cortical primordium. Cereb Cortex 12: 129–139.

Muzio L, Di Benedetto B, Stoykova A, Boncinelli E, Gruss P, Mallamaci A (2002) Conversion of cerebral cortex into basal ganglia in Emx2(−/−) Pax6 (Sey/Sey) double-mutant mice. Nat Neurosci 5: 737–745.

Nitarska J, Smith JG, Sherlock WT, Hillege MM, Nott A, Barshop WD, Vashisht A, Wohlschlegel JA, Mitter R, Riccio A (2016) A functional switch of NuRD chromatin remodeling complex subunits regulates mouse cortical development. Cell Rep 17: 1683–1698.

Ohkubo N, Matsubara E, Yamanouchi J, Akazawa R, Aoto M, Suzuki Y, Sakai I, Abe T, Kiyonari H, Matsuda S, Yasukawa M, Mitsuda N (2014) Abnormal behaviors and developmental disorder of hippocampus in zinc finger protein 521 (ZFP521) mutant mice. PLoS One 9: e92848.

O’Leary DD, Sahara S (2008) Genetic regulation of arealization of the neocortex. Cur Opin Neurobiol 18: 90–100.

Petrich A, Nabhan C (2016) Use of class I histone deacetylase inhibitor romidepsin in combination regimens. Leuk Lymphoma 57: 1755–1765.

Reed GH, Kent JO, Wittwer CT (2007) High-resolution DNA melting analysis for simple and efficient molecular diagnostics. Pharmacogenomics 8: 597–608.

Richards S, Aziz N, Bale S, Bick D, Das S, Gastier-Foster J, Grody WW, Hedge M, Lyon E, Spector E, Voelkerding K, Rehm HL, ACMG Laboratory Quality Assurance Committee (2015). Standards and guidelines for the interpretation of sequence variants: a joint consensus recommendation of the American College of Medical Genetics and Genomics and the Association for Molecular Pathology. Genet Med 17: 405–424.

Sambrook J, and Russell DW (2006) Purification of nucleic acids by extraction with phenol: chloroform. CSH Protoc pdb.prot4455.

Sansom SN, Griffiths DS, Faedo A, Kleinjan DJ, Ruan Y, Smith J, van Heyningen V, Livesey FJ (2009) The level of the transcription factor Pax6 is essential for controlling the balance between neural stem cell self-renewal and neurogenesis. PLoS Genet 5: e1000511.

Saulnier A, Keruzore M, De Clercq S, Bar I, Moers V, Magnani D, Walcher T, Filippis C, Kricha S, Parlier D, Viviani L, Matson CK, Nakagawa Y, Theil T, Götz M, Mallamaci A, Marine JC, Zarkower D, Bellefroid EJ (2013) The doublesex homolog Dmrt5 is required for the development of the caudomedial cerebral cortex in mammals. Cereb Cortex 23: 2552–2567.

Scardigli R, Bäumer N, Gruss P, Guillemot F, Le Roux I (2003) Direct and concentration-dependent regulation of the proneural gene Neurogenin2 by Pax6. Development 130: 3269–3281.

Schaeren-Wiemers N, Gerfin-Moser A (1993) A single protocol to detect transcripts of various types and expression levels in neural tissue and cultured cells: in situ hybridization using digoxigenin-labelled cRNA probes. Histochemistry 100: 431–440.

Scicchitano S, Giordano M, Lucchino V, Montalcini Y, Chiarella E, Aloisio A, Codispoti B, Zoppoli P, Melocchi V, Bianchi F, De Smaele E, Mesuraca m, morrone G, Bond HM (2019) The stem cell-associated transcription co-factor, ZNF521, interacts with GLI1 and GLI2 and enhances the activity of the Sonic hedgehog pathway. Cell Death Dis 10: 715.

Shahbazi E, Moradi S, Nemati S, Satarian L, Basiri M, Gourabi H, Mehrjardi NZ, Günter P, Lampert A, Händler K, Hatay FF, Schmidt D, Molcanyi M, Hescheler J, schultze JL, Saric T, Baharvand H (2016) Conversion of human fibroblasts to stably self-renewing neural stem cells with a single zinc-finger transcription factor. Stem Cell Reports §/ 539–551.

Shen S, Pu J, Lang B, McCaig CD (2011) A zinc finger protein Zfp521 directs neural differentiation and beyond. Stem Cell Res Ther 2: 1–4.

Sobreira N, Schiettecatte F, Valle D, Hamosh A (2015) GeneMatcher: a matching tool for connecting investigators with an interest in the same gene. Hum Mutat 36: 928–930.

Sun J, Rockowitz S, Xie Q, Ashery-Padan R, Zheng D, Cvekl A (2015) Identification of *in vivo* DNA-binding mechanisms of Pax6 and reconstruction of Pax6-dependent gene regulatory networks during forebrain and lens development. Nucleic Acids Res 43: 6827–6846.

Tole S, Hébert J (2020) Telencephalon patterning. In Patterning and cell type specification in the developing CNS and PNS (pp. 23–48). Academic press.

Toresson H, Potter SS, Campbell K (2000) Genetic control of dorsal-ventral identity in the telencephalon: opposing roles for Pax6 and Gsh2. Development 127: 4361–4371.

Urquhart JE, Beaman G, Byers H, Roberts NA, Chervinsky E, O’Sullivan J, Pilz D, Fry A, Williams SG, Bhaskar SS, Khayat M, Simanovsky N, Shachar IB, Shalev SA, Newman WG (2016) DMRTA2 (DMRT5) is mutated in a novel cortical brain malformation. Clin Genet 89: 724–727.

Van Lint C, Ghysdael J, Paras Jr P, Burny A, Verdin E. (1994). A transcptional regulatory element is associated with a nuclease-hypersensitive site in the pol gene of human immunodeficiency virus type 1. J. Virol 68 :2632–2648.

Wilkinson DG, Nieto MA (1993) Detection of messenger RNA by in situ hybridization to tissue sections and whole mounts. Methods Enzymol 225:361–367.

Yelagandula R, Stecher K, Novatchkova M, Michetti L, Michlits G, Wang J, Hofbauer P, Vainorius G, pribitzer C, Isbel L, Mendjan S, Schübeler D, Elling U, Brennecke J, Bell O (2023) ZFP462 safeguards neural lineage specification by targeting G9A/GLP-mediated heterochromatin to silence enhancers. Nat Cell Biol 25: 42–55.

Young FI, Keruzore M, Nan X, Gennet N, Bellefroid EJ, and Li M (2017) The doublesex-related Dmrta2 safeguards neural progenitor maintenance involving transcriptional regulation of Hes1. Proc Natl Acad Sci USA 114: E5599–E5607.

Yousaf H, Fatima A, Ali Z, Baig SM, Toft M, Iqbal Z (2022) A novel nonsense variant in GRM1 causes autosomal recessive spinocerebellar ataxia 13 in a consanguineous Pakistani family. Genes 13: 1667.

Yousaf H, Rehmat S, Jameel M, Ibrahim R, Hashmi SN, Makhdoom EUH, Iwaszkiewicz J, Saadi SM, Rariq M, Baig S, Toft M, Fatima A, Iqbal Z (2023) A homozygous founder variant in PDE2A causes paroxysmal dyskinesia with intellectual disability. Clin Genet 104: 324–333.

Ypsilanti AR, Rubenstein JL (2016) Transcriptional and epigenetic mechanisms of early cortical development: An examination of how Pax6 coordinates cortical development. J Comp Neurol 524: 609–629.

Ypsilanti AR, Pattabiraman K, Catta-Preta R, Golonzhka O, Lindtner S, Tang K, … and Rubenstein JL (2021) Transcriptional network orchestrating regional patterning of cortical progenitors. Proc Natl Acad Sci USA 118: e2024795118.

Yun K, Potter S, Rubenstein JL (2001) Gsh2 and Pax6 play complementary roles in dorsoventral patterning of the mammalian telencephalon. Development 128: 193–205.

Zembrzycki A, Chou SJ, Ashery-Padan R, Stoykova A, O’leary, DD (2013) Sensory cortex limits cortical maps and drives top-down plasticity in thalamocortical circuits. Nat Neurosci 16: 1060–1067.

